# IDSL.UFA assigns high confidence molecular formula annotations for untargeted LC/HRMS datasets in metabolomics and exposomics

**DOI:** 10.1101/2022.02.02.478834

**Authors:** Sadjad Fakouri Baygi, Sanjay K Banerjee, Praloy Chakraborty, Yashwant Kumar, Dinesh Kumar Barupal

**Affiliations:** Department of Environmental Medicine and Public Health, Icahn School of Medicine at Mount Sinai, New York, NY, 10029, USA; Non-communicable Diseases Division, Translational Health Science and Technology Institute, Faridabad, Haryana, 121001, India

## Abstract

Untargeted LC/HRMS assays in metabolomics and exposomics aim to characterize the small molecule chemical space in a biospecimen. To gain maximum biological insights from these datasets, LC/HRMS peaks should be annotated with chemical and functional information including molecular formula, structure, chemical class and metabolic pathways. Among these, molecular formulas may be assigned to LC/HRMS peaks through matching theoretical and observed isotopic profiles (MS1) of the underlying ionized compound. For this, we have developed the Integrated Data Science Laboratory for Metabolomics and Exposomics – United Formula Annotation (IDSL.UFA) R package. In the untargeted metabolomics validation tests, IDSL.UFA assigned 54.31%-85.51% molecular formula for true positive annotations as the top hit, and 90.58%-100% within the top five hits. Molecular formula annotations were also supported by MS/MS data. We have implemented new strategies to 1) generate formula sources and their theoretical isotopic profiles 2) optimize the formula hits ranking for the individual and the aligned peak lists and 3) scale IDSL.UFA-based workflows for studies with larger sample sizes. Annotating the raw data for a publicly available pregnancy metabolome study using IDSL.UFA highlighted hundreds of new pregnancy related compounds, and also suggested presence of chlorinated perfluorotriether alcohols (Cl-PFTrEAs) in human specimens. IDSL.UFA is useful for human metabolomics and exposomics studies where we need to minimize the loss of biological insights in untargeted LC/HRMS datasets. The IDSL.UFA package is available in the R CRAN repository https://cran.r-project.org/package=IDSL.UFA. Detailed documentation and tutorials are also provided at www.ufa.idsl.me.

## Introduction

Untargeted LC/HRMS analyses of human specimens enable studying the metabolome and exposome in an unbiased manner^1, 2^.They have delivered many novel biomarkers and mechanisms for diseases and have improved our understanding of basic metabolic pathways^3-5^. These assays are unique in nature since they record all the mass to charge (m/z) ratio signals above the limit of detection of an instrument for ionized compounds in a sample^6^. This makes the collected data a rich source of information with great opportunities to generate novel hypotheses about metabolome and exposome. It is critical for the promises that untargeted assay offers, that the data are utilized in an inclusive way to not miss any discovery opportunities.

A key post-data acquisition step in the untargeted LC/HRMS assays is to annotate the detected peaks with a range of structural and functional information which can enable biological interpretations^3, 7, 8^. This information includes a chemical structure, molecular formula, chemical class and metabolic pathway^1, 9, 10^. These annotations may help in understanding the nature, origin and function of the chemical structure underlying a peak. Among these information, molecular formula can be assigned to a LC/HRMS peak using the observed and theoretical isotopic profiles for a chemical compound.^11^ Isotopic profiles are distinguishable mass spectral signature that represent atomic masses and their natural abundances in the molecular formulas of a compound.^12^ Despite the known limitations of high-resolution mass spectrometry instruments, observed experimental isotopic profiles for an ionized compound may sufficiently match the theoretical counterpart within instrument errors in many instances^11, 13^, allowing to annotate LC/HRMS peaks with molecular formula^14^. Peak annotation by isotopic profile matching should be performed using efficient computational strategies to account for instrumental errors, multi-sample studies, biological plausibility and chemical diversity.^15^

There has been a great deal of efforts to develop computational tools for annotating peaks in a LC/HRMS dataset with MS1 only data. In a MS1 peak list, a series of m/z values representing different isotopes, ESI adducts, and in-source fragments can belong to one compound. Grouping these m/z values are normally performed by retention time and elution profile similarities within a single file, for example by *xcms*-CAMERA^16, 17^, and peak intensity correlations across multiple samples such as MS-FLO^18^ and CliqueMS^9^ tools. Clustered isotopologues from these tools can be used by the Rdisop R package^19^ to assign molecular formulas in a ‘database independent’ manner. But this approach may miss expected compounds for a sample due to MS instrument’s sensitivity and specificity. MetDNA^7^ can search for theoretical isotope profiles for a list of molecular formulas from a metabolic reaction network database in the MS1 peak list. However, their ‘database dependent’ approach is prone to miss 1) exposure compounds that are poorly represented in such biochemical databases and 2) compounds which may not have any transformation products because of their bioaccumulative nature and 3) compounds that were filtered out by the detection frequency and intensity thresholds while generating MS1 peak table for a study. Moreover, MetDNA^7^ and other tools including SIRIUS^20^ and ZODIAC^21^, NetID^22^ are mainly designed for assigning molecular formulas to peaks having MS/MS fragmentation data. Furthermore, implementing these tools for larger studies where only MS1 data are available for every sample remains to be challenging due to the ranking of formula hits on individual and aligned peak tables, scalable computation and various sources for formulas which need to be covered for exposomics projects.

There is a need to develop new tools to compute and to compare theoretical and experimental isotopic profiles for chemical lists from larger databases and chemical spaces for molecular formula annotations. Here, we have developed a scalable, user-friendly, thoroughly tested R package, the IDSL.UFA to assign molecular formulas with high confidence to peaks in untargeted LC/HRMS datasets from large-scale studies. IDSL.UFA covers major possible situations in which a molecular formula can be assigned to LC/HRMS peaks. We propose that processing LC/HRMS data with IDSL.UFA can find new opportunities for hypotheses and biomarker discoveries for studying the role of metabolism and exposome in human diseases.

## Methods

### Publicly available LC/HRMS test datasets

To test and develop the IDSL.UFA R package, we have utilized the raw LC/HRMS data for human and mouse biospecimen studies (MTBLS1684^23^, MTBLS2542^24^, ST001683^25^, ST001430^26^, ST001154^27^, ST002044 and reference authentic standards (MSV000088661) available from Metabolomics WorkBench (https://www.metabolomicsworkbench.org/), MassIVE (https://www.massive.ucsd.edu), and MetaboLights (https://www.ebi.ac.uk/metabolights) repositories. Data processing results that we have generated for these studies have been submitted to the Zenodo.org repository and corresponding entry pages are provided in Table S.1. Sample preparation and data collection procedures are available at entry pages for these studies in the repositories.

### Data analysis setup

IDSL.UFA R package is available in the R-CRAN repository (https://cran.r-project.org/package=IDSL.UFA). The package was installed using the ‘*install*.*packages(“IDSL*.*UFA”)’* R command. The IDSL.MXP package (https://cran.r-project.org/package=IDSL.MXP) was used to read mzML/mzXML/netCDF mass spectrometry data in the centroid mode. mzML files were generated from the vendor specific data format using the ProteoWizard MSConvert utility^28^ when needed. All data files related to only one type of analysis such as “reverse phase - electrospray ionization negative mode” were stored in a single file folder. Figure 1 (simplified) and Figure S.1 (detailed) show the workflow steps to assign molecular formulas for a study. Data processing parameters for IDSL.UFA were provided in a Microsoft excel file (https://zenodo.org/record/6466688) which was created for individual test studies. We have provided the parameters files used in this manuscript in the Zenodo.org repository at (https://zenodo.org/record/6466684). To run the IDSL.UFA workflow, only a single R command ‘*UFA_workflow*(*spreadsheet = “address of the parameter xlsx file”*)’ was needed. Tutorials to create the parameter files for different scenarios are available at (https://ufa.idsl.me). For each individual peak list in a study, a formula annotation list with rank, score and other peak properties was generated and exported to a csv file. Likewise, for each peak in the aligned peak table, top 5-20 formulas with detection frequency and median ranks across all samples were exported to a csv file for each test study.

**Figure 1.**
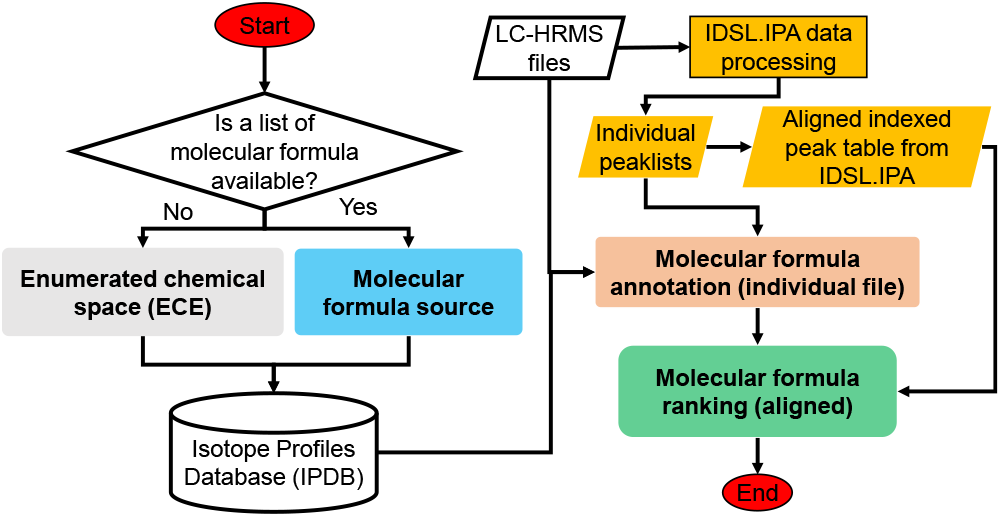
A simplified flowchart of the IDSL.UFA software.

### Generating the isotopic profile database (IPDB)

An IPDB is a digital collection of theoretical isotopic profiles computed by the IDSL.UFA R package for a list of candidate molecular formulas. IDSL.UFA queries and matches the experimental isotopic profile against this collection to annotate a LC/HRMS peak. To compute the isotopic profile for a molecular formula, we have utilized the reference stable isotope masses and abundances for elements in the periodic table from the PubChem database entries^29^ which have been sourced from International Union of Pure and Applied Chemistry (IUPAC)^30^. We have also provided an online tool (https://.ipc.idsl.me) to compute an isotopic profile for a single molecular formula. IDSL.UFA generates centroid isotopic profiles using a dynamic intensity threshold and a peak-spacing criterion to merge adjacent isotopologues within a mass accuracy window. In this work, we have covered two sources of molecular formulas.

### Source A (databases)

Chemical compound lists for four key databases in metabolomics and exposomics including the blood exposome (chemicals expected in a mammalian blood specimen), RefMet (measured and expected small molecules in biological organisms)^31^, Lipid Maps (known lipid molecules)^32^ and the US-Food and Drug Administration substance registry^33^ were obtained from their online web addresses. These four databases were combined into a single compound list referenced as IDSL.ExposomeDB in this manuscript and also provided at the Zenodo repository (https://zenodo.org/record/5823455). Charged compounds, isotope-labeled compounds and multi-components were excluded. Unique molecular formulas from this consolidated database were used for computing IPDB. IPDBs for these four databases and the environmental protection agency (EPA) CompTox Chemicals Dashboard^34^ are available at the Zenodo repository (https://zenodo.org/record/5823455).

### Source B (enumerated chemical space with constraints)

Molecular formulas were enumerated using a set of combinatorial and filtering rules using C, H, As, B, Br, Cl, F, I, K, N, Na, O, P, S, Se, and Si elements. These 16 elements were able to cover 93.76% of carbon-containing compounds (50 ≤ mass ≤ 2000) in the IDSL.ExposomeDB combined with EPA chemistry Dashboard^34^. An enumerated chemical space (ECS) can be represented using equation (1).

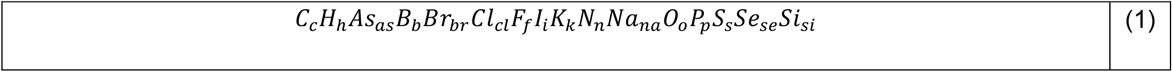

where the subscripts of elements represent the number of atoms. A fully combinatorial chemical space from above-mentioned 16 elements is impractical to be managed by current computational resources. Therefore, we derived and coded in R a set of four rules which were inspired from the seven golden rules approach^35^ to constrain ECSs. These rules included **1) C/N chemical space rule** ‘((c/2-n-1) ≤ (h+cl+br+f+i) ≤ (2c+3n+6))’ was used to set elemental boundaries for the organic compounds to ensure entire moieties are bond to carbon and nitrogen atoms. **2) Extended SENIOR rule** was used to ensure that the molecular formulas completely filled s- and p-valence electron shells.^35^ **3) Maximum number of halogens thresholds** was used to constrain halogenated compounds. For example, we have used the maximum number of (br+cl) ≤ 8 and the maximum number of ((br+cl+f+i) ≤ 31) thresholds to cover halogenated compounds in the blood exposome database. **4) Maximum number of elements rule** was used to skip unrealistically complex molecular formulas generated through molecular formula enumeration. For example, the maximum number of elements for glucose (C_6_H_12_O_6_) is three (C, H, and O). The ECS boundaries and rules for the MTBLS1684 study are provided in the Zenodo repository (https://zenodo.org/record/5838603).

### MS1 peak detection and alignment

IDSL.IPA^36^ R package (https://cran.r-project.org/package=IDSL.IPA) was used to generate individual peak lists for each sample and the aligned peak table (m/z-RT pairs across all samples) for each study. Data processing parameter files and IDSL.IPA results for each test study are provided in the Zenodo repository (see Table S.1). Details and a tutorial for IDSL.IPA data processing can be found at (https://ipa.idsl.me) site.

### Isotopic profile matching for individual sample

First, IDSL.UFA software accessed the peak boundaries, ^12^C m/z, ^13^C m/z and ratio of cumulated intensity of ^12^C to ^13^C (R^13^C) for each peak in an IDSL.IPA generated peak list for a sample. Next, it finds all the theoretical isotopic profiles in an IPDB that matches the ^12^C and ^13^C m/z for a peak. Then, for each matched theoretical profile, experimental profiles are retrieved from raw data using a mass accuracy threshold within the peak boundaries for a peak. If a compound formula has three isotopologues in the IPDB and only two were observed in the raw data, the formula will not be annotated. IDSL.UFA requires that a minimum one MS1 scan across the peak should have the full isotope profile for a formula in the IPDB.

For the experimental isotopic profiles, the IDSL.UFA software calculates cumulated intensities and intensity-weighted average masses for each isotopologue using equations (2) and (3) across the chromatographic peak to minimize the effect of fluctuations such as peak saturation.

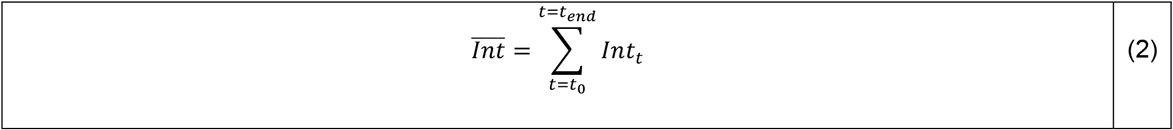

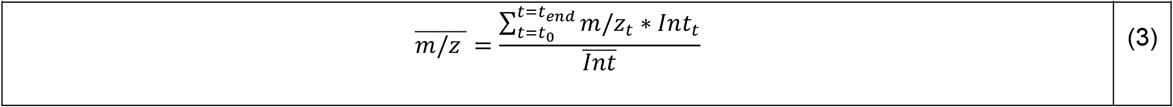

where *m/z*_*t*_ and *Int*_*t*_ represent mass and intensity of the matched isotopologue in individual scans across the chromatographic peak from *t*_*0*_ to *t*_*end*_.

We have used the Profile cosine similarity 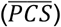 to quantify profile similarity between experimental and theoretical isotopic profiles using equation (4). To assess mass accuracy error for whole isotopic profile, Normalized Euclidean mass error 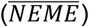 was calculated using the equation (5).^11^

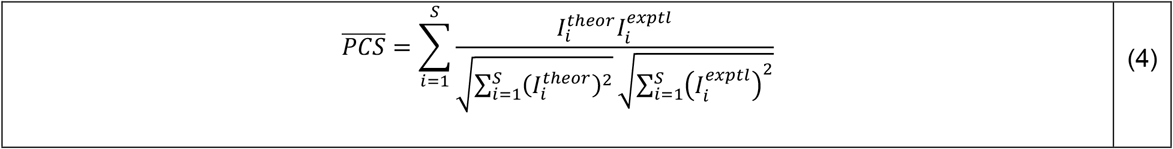

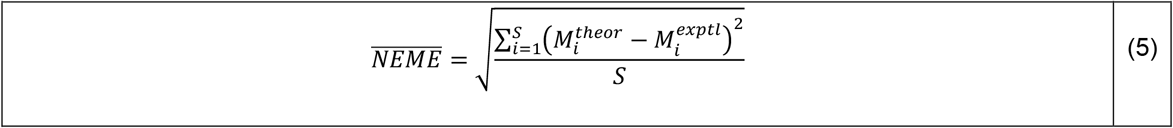

where *I*_*i*_, *M*_*i*_, and *S* represent the intensity of the isotopologue, mass of the isotopologues, and number of isotopologues in the isotopic profile, respectively. Superscripts of *theor* and *exptl* also represent theoretical and experimental isotopic profiles, respectively.

Candidate formulas were then filtered using thresholds for 1) 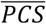 2) 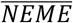 3) the top 80% of number of scans with the confirmed whole isotopic profile (NDCS) and 4) minimum percentage of NDCS within a chromatography peak (RCS (%)). These linear cutoffs allow eliminating false positives; however, they can reject true positive peaks with poor isotopic profiles.

Next, a matching score for each candidate filtered formula was computed using equation (6).

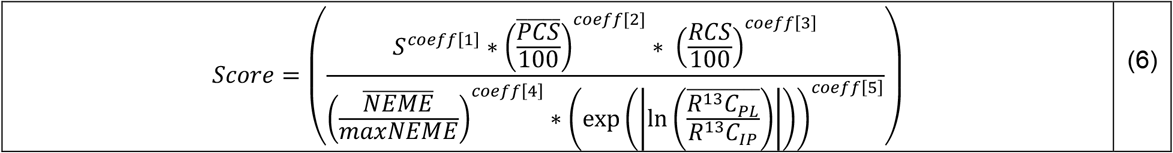

where 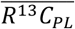 and *R*^13^*C*_*IP*_ indicate experimental and theoretical R^13^C values, respectively. R^13^C values represent the ratio of the general ^13^C isotopologue [M+1] relative to ^12^C isotopologue [M] on the most abundant mass. *coeff[1-5]* are powers of the parameters to apply different magnitudes of each variable in different studies. Using this score, a ranking for candidate formula was determined. By default, IDSL.UFA utilized a value of 1 for *coeff[1-5]* to rank candidate molecular formulas in the equation (6). However, we have provided a score coefficient optimization strategy in the section S.1 which can be helpful for improving the ranking when larger size IPDB are utilized.

### Summary statistics of molecular formulas annotation in the aligned peak table

It is quite common to have more than 50 samples in metabolomics and exposomics projects, which can be leveraged to compute a statistic for formula annotations across all the samples. For each peak (*m/z*-RT pair) in the aligned peak table, corresponding molecular formula lists across all the samples were retrieved using the peak indices provided by the IDSL.IPA data processing. We then aggregated these formula lists and computed two properties 1) the detection frequency and 2) median rank for each formula assigned for a peak across all the samples (individual peak list). Then we generated a new sort order for each molecular formula at the aligned peak table level using the following formula: 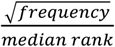. For each peak in the aligned peak table, top 5-20 formulas with detection frequency and median ranks across all samples were exported to a csv file for each test study.

### Molecular formula class detection

Many compounds belong to a chemical class with a distinct sub-structure pattern such as polychlorinated biphenyl (PCBs), polybrominated diphenyl ethers (PBDEs), polycyclic aromatic hydrocarbons (PAHs), perfluoroalkyl substances (PFAS), lipids and phthalates etc. The formula annotations generated via the enumerated chemical space (ECS) approach were processed to detect such classes within a list of formulas. The IDSL.UFA function ‘*detect_formula_sets*’ was used to detect 1) constant ΔH/ΔC ratios for polymeric (ΔH/ΔC = 2) and cyclic (ΔH/ΔC = 1/2) chain progressions within polymeric and cyclic classes (Table S.2-S.4) and 2) a constant number of carbons and fixed summation of hydrogens and halogens (Σ(H+Br+Cl+F+I)) representing classes similar to PCBs, PBDEs (Table S.5).

### Correlation analysis for gestational age

The ST001430 study^26^ includes weekly blood samples of 30 pregnancies. The study has 781 total samples each processed in positive and negative modes to predict gestational age. To reduce batch effects, the peak heights were adjusted by raw total ion chromatograms (TICs) in each sample, and then the positive and negative aligned peak height tables were stacked to generate a comprehensive list of peaks. We computed a Spearman correlation coefficient between gestational age and peak height data for each pregnancy. A schematic of this workflow is presented in Figure S.2.

## Results and discussion

We have engineered a new software, IDSL.UFA, to annotate LC/HRMS peaks with molecular formulas for an untargeted metabolomics or exposomics study. In this approach, IDSL.UFA computes theoretical isotopic profiles for molecular formulas, matches theoretical isotopic profiles against experimental LC/HRMS data in individual data file using a set of matching parameters and then summarizes the formula annotations using detection frequency and median ranks in multiple samples (aligned annotated peak table) in a study. The IDSL.UFA software has been implemented as an R package and made publicly available via the R-CRAN repository and www.ufa.idsl.me site.

### Section 1) Development and validation of IDSL.UFA results

To demonstrate the validity of our approach to assign molecular formulas, we have utilized datasets with true positive annotations and show their ranks in the IDSL.UFA result matrices.

### Analysis of authentic reference standards

First, we evaluated performance of the IDSL.UFA software to detect molecular formulas in LC/HRMS data for authentic reference standards. We found that the average 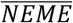 (indicator of mass difference) was 0.70 mDa and 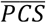 (indicator of isotope profile similarity) were 99.968% between experimental and theoretical isotopic profiles for 367 authentic standard compounds of common metabolites. This indicated that the observed isotopic profiles were very similar to the theoretical counterparts for these reference standards and suggested that molecular formulas can be reliably assigned to untargeted data generated by the commonly used LC/HRMS instruments. The theoretical and experimental integrated isotopic profile spectra across chromatography for these standards are provided at Zenodo repository accession (https://zenodo.org/record/5803968) and an example compound (Kynurenine ion [C_10_H_13_N_2_O3]^+^) is shown in Figure 2 and Figure S.3 (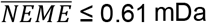 and 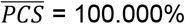).

**Figure 2.**
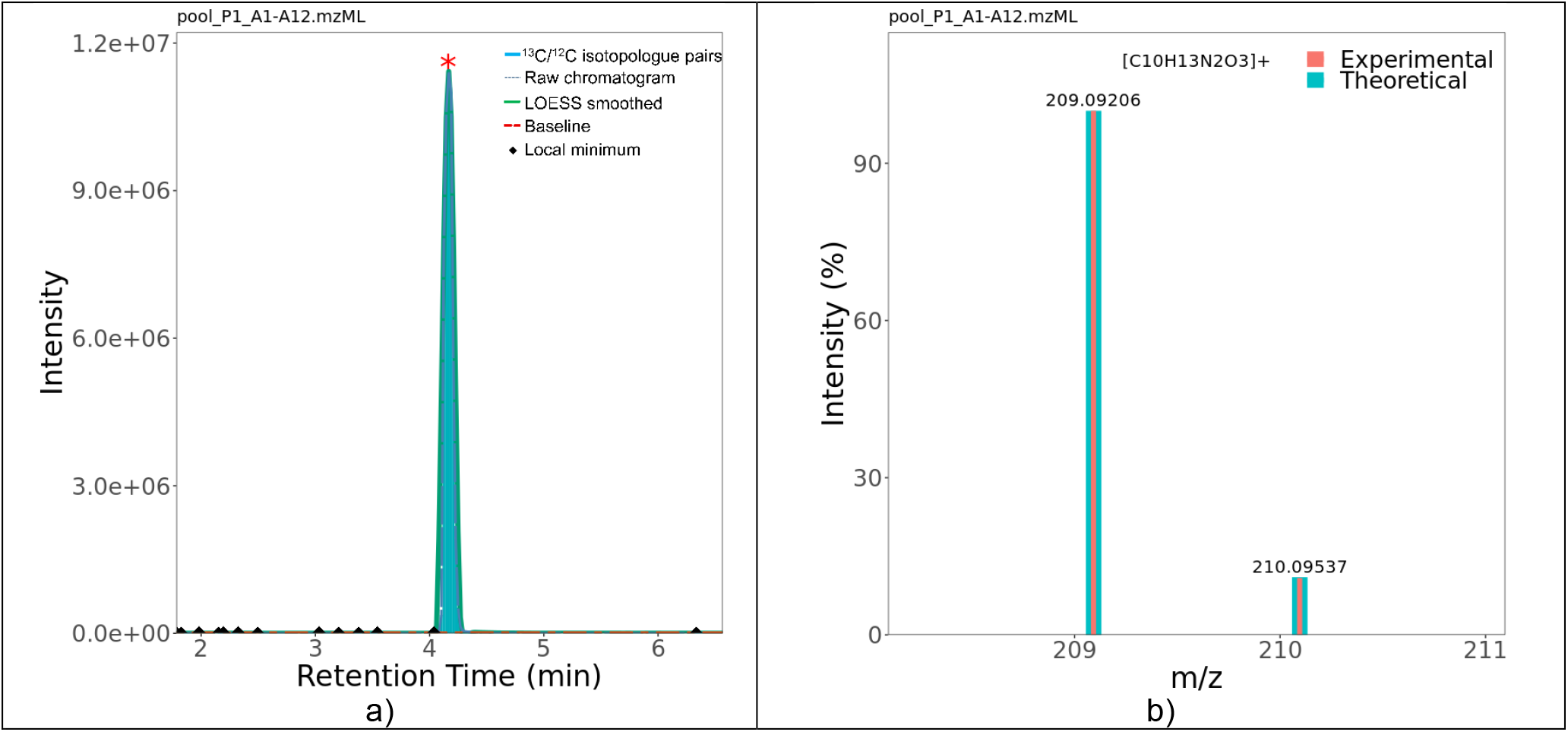
a) A chromatographic peak generated by the IDSL.IPA pipeline for Kynurenine ion ([C_10_H_13_N_2_O_3_]^+^ = [M+H]^+^) to detect peak boundaries. b) Comparison between the theoretical isotopic profile and integrated spectra across the chromatographic peak after molecular formula annotation using IDSL.UFA.

### Analysis of untargeted LC/HRMS data with structurally annotated peaks

We selected four publicly available studies (ST001154^27^, ST001683^25^, MTBLS1684^23^, and MTBLS2542^24^). These studies have reported annotations with MSI 1-3 confidence levels (https://zenodo.org/record/5838709) that were obtained using retention time, accurate mass and MS/MS spectra matching. For these studies, the IDSL.UFA software assigned 61.85%, 54.31%, 70.58% and 85.51% molecular formula as the top hit, and 96.90%, 90.58%, 100% and 99.29% molecular formulas in the top five hits in the aligned table. These results were generated using the IPDB of the IDSL.ExposomeDB with 209,592 and 129,122 ion formulas in positive and negative modes from multiple ionization pathways, respectively representing 83,951 unique intact molecular formulas (http://zenodo.org/deposit/5838709).

For each selected study, an ECS IPDB was generated using the element boundaries that covered the formula list of true positive annotations for the study. When IDSL.UFA software was used for each study using those specific ECS IPDBs, the assignment rates were – 52.74%, 53.36%, 79.41% and 51.08% molecular formula as the top hit, and 95.60%, 84.45%, 100% and 91.66% molecular formula in the top 5 hits in the aligned table (Figure S.4). Generally, the IDSL.UFA software annotated 924 (90.14%) and 877 (85.56%) molecular formulas across all four studies using IDSL.ExposomeDB and ECS IPDBs, respectively. These findings demonstrate that IDSL.UFA is a sensitive approach to cover the majority of formulas for chemicals detectable in a biospecimen.

There is a tradeoff of coverage and the confidence in annotation while choosing chemical space for molecular formula annotation. We have noticed that the rank of true positive hits degrades when we have used a larger chemical space (Figure S.5). However, when compounds that are known and expected to be found in a blood specimen are used, we have observed that formulas for true positives are often ranked top hits. Therefore, we recommend a chemical prioritization strategy by sample type and to first match the compounds that are expected for that sample type and then expand the chemical space to cover additional peaks.

### Summary of the formula annotations in the aligned peak table

Our raw data processing generates both a separate list of m/z-RT pairs for each sample (individual peak list) and a single combine list (aligned-table) of m/z-RT pairs for all samples. IDSL.UFA annotates molecular formulas only to individual peak lists, then, it computes the detection frequency and median rank for all formulas annotated for the same peak across all samples using the aligned peak table (See methods). Our hypothesis is that the most probable formula of the underlying ionized compound will have a higher detection frequency and median rank across all the samples. For example, for the MTLS1684 study, 24/35 (69%) of the reported annotations had a median rank of 1 and 8/35 (23%) had a median rank of 2 across all 499 samples (https://zenodo.org/record/5838709). We propose that the summary of detection frequencies and ranks across individual data files can be helpful in boosting the confidence for formula assignments in multi-sample studies. It should be noted that IDSL.UFA does not group related peaks to flag them as potential ESI adducts or in-source fragments. Such grouping of peaks can be achieved by existing solutions such as MS-FLO^18^ online tool or CliqueMS^9^ R package.

### Additional validation of molecular formula assignment by MS/MS

To further ensure that IDSL.UFA can assign high confidence molecular formulas for untargeted LC/HRMS data, we utilized data from ST002044 study which has high quality MS/MS data collected in the data dependent mode. A total 73 hits were confirmed by matching their spectra to the NIST 2020 MS/MS library (https://chemdata.nist.gov) and public mass spectral libraries (https://zenodo.org/record/6416108). IDSL.UFA assigned 78.75% of these hits within a median rank of ≤ 2 in the aligned peak table generated using the IDSL.ExposomeDB formula IPDB (Table S.6 and Figure S.6). These results provided additional supports to confidence in the molecular formula assignment by the IDSL.UFA software using the IDSL.ExposomDB IPDB.

### Rank score optimization

IDSL.UFA utilized a number of chromatographic-mass spectrometry parameters to compute the rank of a molecular formula for a peak in the individual peak list. By default, a score coefficient of 1 is used which works sufficiently in most situations. However, the rank can be further improved by an optimization strategy that utilizes the true positive, curated and high-quality structure annotations for each data file as input. This can be achieved by running a mixture of reference standards using the same analytical method or by annotating peaks using MS/MS, RT and isotopic profile matching using stringent criteria. For metabolite standards (MSV000088661) and blood specimens (ST002044) studies, we have observed a significant improvement in the ranking of molecular formulas when optimized score coefficients were utilized in the IDSL.UFA software (Table S.7).

### Section 2) Application of IDSL.UFA for a pregnancy study

To demonstrate an application of IDSL.UFA software to characterize the metabolome and exposome for blood specimens, we have re-processed a publicly available study ST001430^26^ (n=781) which has weekly blood samples analyzed for 30 pregnancies to accurately predict gestational age (GA in weeks). Raw data were processed using the IDSL.IPA software to generate the individual peak lists and the aligned peak table (https://zenodo.org/record/5804527). On average, (3,416 ESI^-^ and 6,978 ESI^+^) peaks were detected across individual peak lists for this study and a total of (89,174 ESI^-^ and 143,712 ESI^+^) peaks were reported in the aligned peak table. The IDSL.UFA software using the IDSL.ExposomeDB IPDB annotated (80,957 ESI^-^ and 124,647 ESI^+^) peaks in the aligned peak table with at least one molecular formula having a median rank of ≤ 5.

We identify the peaks that were associated with GA by computing a spearman correlation coefficient between normalized peak-height for each peak and GA. On a spearman cutoff of (*p*-value ≤ 0.05, |*ρ*| ≥ 0.65, “two.sided” alternative), 274 peaks with a detection frequency of ≥ 5 within each subject were found to be significantly associated with GA (only ≤ 36 weeks). We observed 242 (red) and 32 (blue) ascending and descending correlations patterns with GA, which were consistent with the patterns reported in the original paper^26^ and corresponded to chemicals related to steroid hormone biosynthesis and long-chain fatty acids. These results show the potential the IDSL.UFA approach to characterize the pregnancy related metabolic changes (Figure 3).

**Figure 3.**
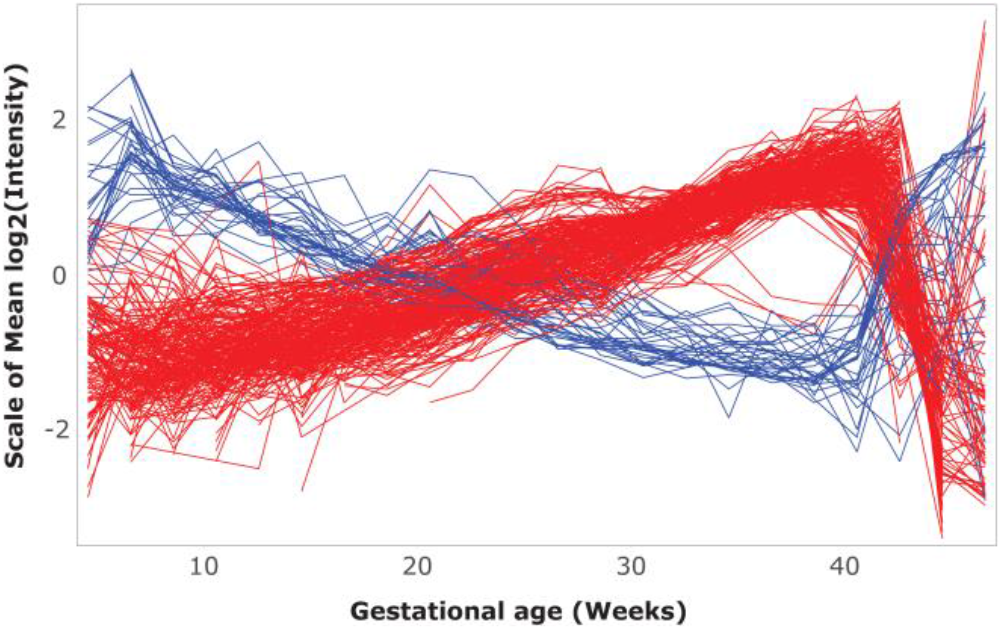
Trends of 274 peaks associated with pregnancy dynamics. Molecular formula annotated interactive plots are available at https://ufa.idsl.me/st001430 for an enumerated chemical space and IDSL.ExposomeDB IPDBs.

To flag the potential peaks related to chemical exposures in the pregnancy study (ST001430), we first assigned a molecular formula using an ECS that may cover diverse halogenated compounds that were not found in the IDSL.ExposomDB formula list. IDSL.UFA resulted with 199,837 unique molecular formulas on the aligned table (top rank ≤ 30 and number of hits ≤ 30) in the ST001430 study. Grouping these formulas by a class detection approach (see method) highlighted that 7,615, 18,452, and 32,107 distinct formula classes. For instance, a class of heavily halogenated compounds, C_n_HClF_2n_O_4_ (n=10-12), known as chlorinated perfluorotriether alcohols (Cl-PFTrEAs) was detected for human specimens in this study. Cl-PFTrEAs was previously only reported in air samples from eastern China^37^ and may represent a new ubiquitous global contaminant class. IDSL.UFA can only confirm isotopic profiles match (Figure S.6); however, a confirmatory in-source fragment ([M-C_3_F_6_O]^-^) was consistent with the published MS/MS fragmentation (Figure S.8).^37^ Authentic standards for Cl-PFTrEAs are not readily available; therefore a confidence level 3b (isotopic profile match combined with fragmentation-based candidate) is suggested for these annotations according to a recently proposed PFAS identification confidence level by Charbonnet et al.^38^ Levels of Cl-PFTrEAs were similar to the commonly known legacy halogenated compounds^14^ for human serum samples (Figure 4). These findings also show that IDSL.UFA software can potentially detect chemicals of public health concerns in a human biospecimen and can be helpful in expanding the existing database of exposome chemicals.^39^

**Figure 4.**
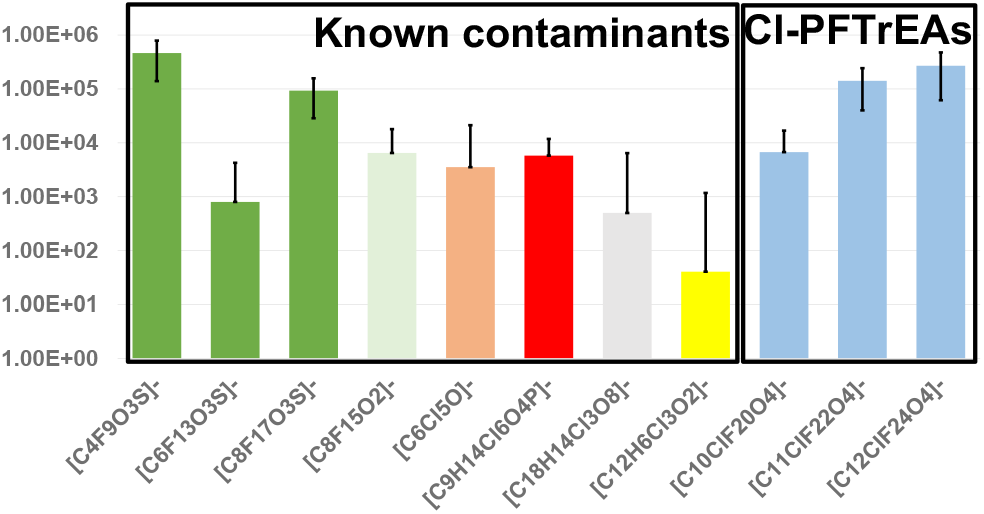
Peak area of halogenated contaminants in human blood ([C_n_F_2n+1_O_3_S]^-^ (n = 4, 6, 8), [C_8_F_15_O_2_]^-^, [C_6_Cl_5_O]^-^, [C_9_H_14_Cl_6_O_4_P]^-^, [C_18_H_14_Cl_3_O_8_]^-^, [C_12_H_6_Cl_3_O_2_]^-^) and Cl-PFTrEAs [C_n_ClF_2n_O_4_]^-^ (n = 10-12) across 781 negative samples in the ST001430 study.

### Section 3) Performance benchmarking and comparison with existing tools

IDSL.UFA processed one file (D115_NEG.mzml from the ST2044 study) in ∼10 minutes on a computer with 6 cores, indicating the pipeline can be used in normally available computing resources.

To check how IDSL.UFA performed for low abundant signals, we utilized data from the MTBLS1040 study which has a seven-point calibration curve for the analyzed compounds. For the hippuric acid standard in the MTBLS1040 study, IDSL.UFA correctly assigned the molecular formula to the corresponding peak in samples analyzed at up to 8 fmol concentration level (second-lowest point) (https://zenodo.org/record/6466668).

IDSL.UFA software is designed to cover commonly used LC-HRMS instruments for human biospecimens studies in the EBI MetaboLights and Metabolomics Workbench repositories. A mass resolution of 20,000 and mass accuracy of 5 ppm is often found for these instruments. We compared the results for publicly available two raw data files for a BioRec human plasma sample analyzed for a lipidomic assay by QToF(ST001843) and Orbitrap instruments(ST001264) using the same chromatography method in the same lab. Our workflow generated 1752 peaks with 2855 formulas for the QToF data file and 1328 peaks with 2209 formulas for the Orbitrap data file. A list of 151 true positive annotations from the ST110054^27^ study (MS/MS matches were inspected by an expert user from the same lab and chromatography method) was utilized for these test data files (https://zenodo.org/record/6621138). For these true positives, 35% were found to be top hits in the QToF data file and 53% in the Orbitrap data file. It seems our approach works slightly better for Orbitrap data. However, an even higher resolution and better mass accuracy can be helpful in removing several false positive annotations, and in improving the ranking of the true positive annotations.

When we imported a MS1 only data file in the SIRIUS^20^ tool, it did not process the file, which was expected since SIRIUS only processes data files with MS/MS spectra. For a data file (D115_NEG.mzml from the ST002044 study) with MS/MS spectra in the Data Dependent Acquisition (DDA) mode, SIRIUS processed 885 MS/MS spectra and suggested formula annotations for 221 spectra, whereas IDSL.UFA assigned molecular formula to 9303 peaks in this data file.

IDSL.UFA natively uses IUPAC isotope table data^29^ to calculate theoretical isotopic profiles and calculated almost identical isotopic profiles to that obtained from the *envi*Pat package^40^ (Table S.8). Negligible mass and profile similarity differences (NEME ≤ 0.69 mDa and PCS ≥ 99.999%) were observed for formula [C_8_F_17_O_3_S]^-^ between IDSL.UFA and *envi*Pat^40^.

Next, we compared the IDSL.UFA against Rdisop^19^ R package to show the advantages of a database-dependent approach (IDSL.UFA) over a database independent approach (Rdisop) for molecular formula annotation. For kynurenine authentic standard (MSV000088661), both IDSL.UFA and Rdisop^19^ ranked the M+H adduct formula as the top hit(Section S.2 and Table S.9). But Rdisop‘s ranking for PFOS isomers were >20 in the studies ST001430 and ST002044 (both human blood samples). Whereas IDSL.UFA annotated both isomers of PFOS as top hit for these studies (Table S.10-11 and Figure S.9-10). This suggests that Rdisop may miss important expected compounds when a complex chemical space (CHBrClFNOPS) is targeted, but IDSL.UFA will be able to annotate them for human blood specimens. Next, we extended the comparison to the lipidomics analysis with 151 true positive annotations. Rdisop annotated 12%, whereas IDSL.UFA reported 53% of true annotations as top hits for the Orbitrap data file (Figure S.11). These comparisons suggest that a database dependent approach for formula annotation, such as IDSL.UFA should be used first to screen for expected compounds in HRMS data before looking for unknown-unknowns. We also provide a comparison (Table S.12) between IDSL.UFA and Rdisop^19^ R packages, highlighting new features that IDSL.UFA is introducing into R computing workflows for metabolomics and exposomics studies.

Our approach to obtain homologous series with polymeric chain increment from a list of input molecular formulas is different from the prior approaches^41-43-40^ in which molecular formulas are enumerated only for a known series or chain increment rule. Therefore, our approach has the flexibility to discover new types of homologous series among a collection of formulas.

## Conclusion

IDSL.UFA enabled a comprehensive characterization of the chemical space that was detected by an untargeted LC/HRMS assay to study the metabolome and exposome and its role in human health. The unique feature of the IDSL.UFA software is to utilize the summary statistics for the rank and frequency of detected molecular formulas in the aligned annotated molecular formula table. It can complement the other peak annotation efforts that use mainly MS/MS data to annotate peaks, and thus lower the number of false negative reporting of peaks and minimize the under-utilization of the untargeted LC/HRMS datasets. We provided various scenarios to obtain molecular formulas from a known database and enumeration strategies to assign a formula to peaks in a LC/HRMS dataset. These new computational strategies for molecular formula assignment can greatly expand the quality of untargeted LC/HRMS data matrices and their analyses especially when MS/MS data are not available.

## Author’s contribution

SFB and DKB planned the study, prepared the results and drafted the manuscript. SFB and DKB coded the IDSL.UFA package. YK, SB and PC provided test LC/HRMS data for authentic standards of metabolites and human biospecimens. All authors have reviewed the manuscript content.

