## Supplemental Information for "IDSL.UFA assigns high confidence molecular formula annotations for untargeted LC/HRMS datasets in metabolomics and exposomics"

**Table of contents**

**Figure S.1.** A detailed flowchart of the IDSL.UFA software. (*MAIso represents the most abundant isotopologue)

**Table S.1.** LC/HRMS datasets used in this work

**S.1.** Objective functions for the score coefficients optimization

**Table S.2.** Polymeric carbon chain progression for PFAS

**Table S.3.** Cyclic chain progression for polycyclic phenols

**Table S.4.** Repeated linear and cyclic groups in a class of compounds

**Table S.5.** Constant numbers of carbons and Σ(H+Cl) for PCBs

**Figure S.2.** Schematic of workflow to annotate peaks with linear patterns in pregnancy.

**Figure S.3.** Comparison between theoretical and integrated experimental isotopic profiles across a chromatographic peak after molecular formula annotation using the IDSL.UFA pipeline on an authentic standard from the MSV000088661 study.

**Figure S.4.** Distribution of the candidate order on the aligned annotated molecular formula tables in each study. (* IPDBs may be found at <https://zenodo.org/record/5823455>)

**Figure S.5.** Evaluation of performance of the ranking method for true positives (only [M+H]^+^ ionization used) with respect to various chemical spaces on the MSV000088661 study (size of chemical space is shown inside parenthesis). (zenedo address)

**Table S.6.** Comparison between aligned annotated table using IDSL.ExposomeDB IPDB by the IDSL.UFA and MS/MS library match for the ST002044 study.

**Figure S.6.** Distribution of the candidate order on the aligned annotated molecular formula table using IDSL.ExposomeDB IPDB for the matched compounds using the MS/MS library shown on Table S.7 for the ST002044 study. A set of relaxed criteria (generally mass accuracy ≤ 7.5 mDa, $\bar{NEME}$ ≤ 7.5 mDa, $\bar{PCS}$ ≥ 95%) was used for this study.

**Table S.7.** Evaluation of performances of the objective functions on different datasets

**Figure S.7.** Examples of the chromatographic peaks and integrated mass spectra pertaining to the Cl-PFTrEAs ([C*_n_*ClF_2_*_n_*O_4_]^-^ (where *n* = 10-12) class in the ST001430 study.

**Figure S.8.** A confirmatory fragment ([M-C_3_F_6_O]^-^) pertaining to [C_12_ClF_24_O_4_]^-^ present in MS1 from the ST001430 study and also present in the published MS/MS spectra. Mass spectra were collected from apexes of peaks using MZmine 2.0^1^.

**Table S.8.** Comparison between calculated isotopic profile of [C_8_F_17_O_3_S]^-^ by *envi*Pat^2^ and IDSL.UFA

**S.2.** Rdisop command details

**Table S.9.** Comparison of ranks of two adducts of Kynurenine (C_10_H_12_N_2_O_3_) in ”pool_P1_A1-A12.mzML” from MSV000088661 study among three different annotation methods on a simple chemical space

**Table S.10.** Comparison of ranks of two isomers of PFOS ([C_8_F_17_O_3_S]^-^) in”1.NEG.mzXML” from ST001430 study between annotation methods on a complex chemical space ( CHBrClFNOPS)

**Figure S.9.** EIC and integrated spectra of the PFOS peaks in 1.NEG.mzXML” from ST001430

**Table S.11.** Comparison of ranks of PFOS (C_8_HF_17_O_3_S) in ”D_194NEG.mzML” from ST002044 study between annotation methods on a complex chemical space

**Figure S.10.** EIC and integrated spectra of the PFOS peak in ”D_194NEG.mzML” from ST002044 study

**Figure S.11**. **Figure S.11.** Comparison of ranks of 151 true positive hits between IDSL.UFA and Rdisop.

**Table S.12.** Summarized operational comparison between IDSL.UFA and Rdisop^3^.

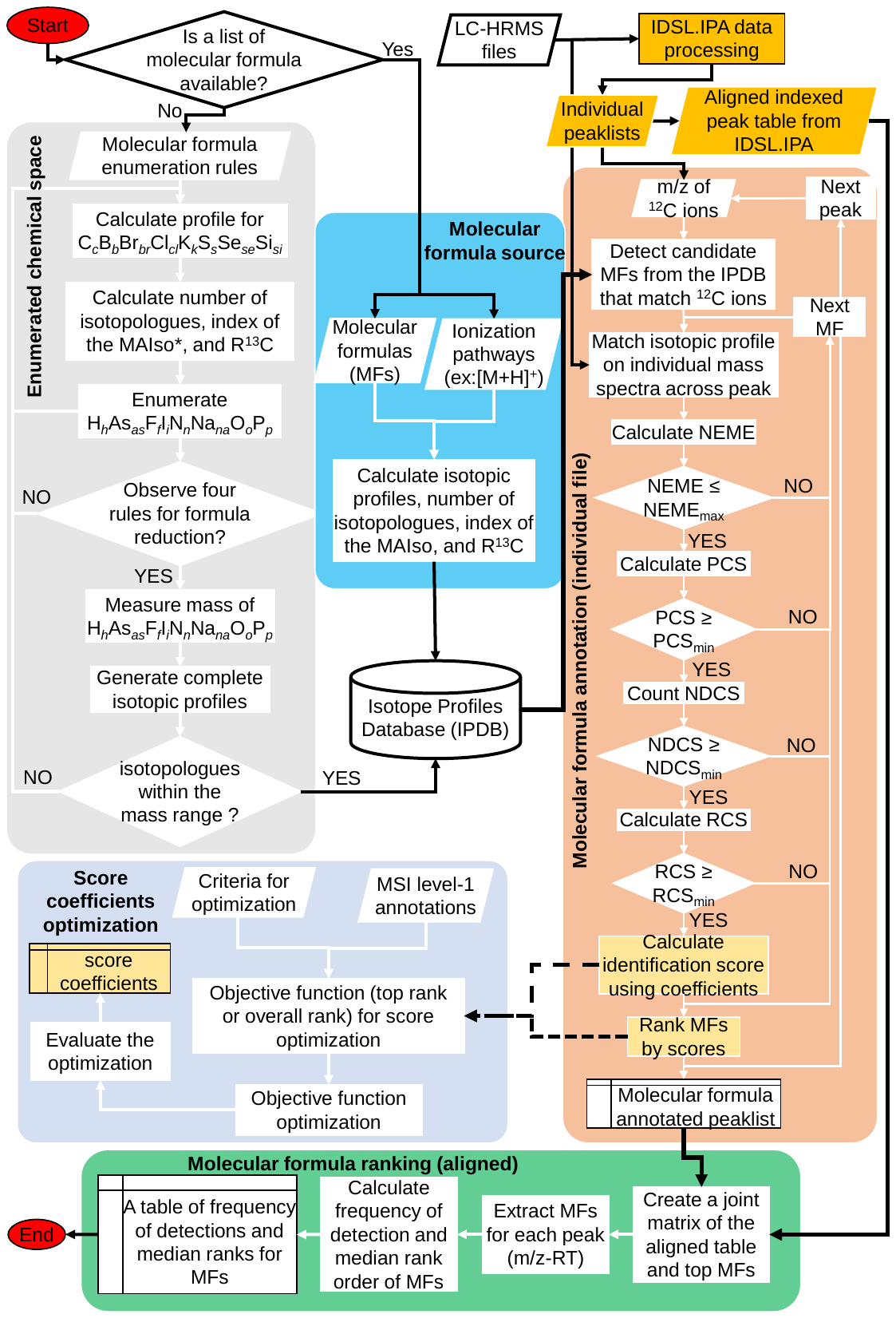

**Figure S.1.** A detailed flowchart of the IDSL.UFA software. (*MAIso represents the most abundant isotopologue)

| **Table S.1.** LC/HRMS datasets used in this work | | | | | | | | |
| --- | --- | --- | --- | --- | --- | --- | --- | --- |
| **Sample Type** | **Repository *** | **Accession ID** | **Analysis mode** | **Mass analyzer** | **Used mass accuracy (mDa)** | **Number of samples** | **Total number of compounds** | **IDSL.UFA and IDSL.IPA results DOI (Zenodo.org)** |
| Cord blood | EBI-Metabolights | MTBLS1684^4^ | RP – ESI – POS | Agilent 6550 QTOF | 10 | 499 | 34 | <https://zenodo.org/record/5803978> |
| Blood (COVID-19) | EBI-Metabolights | MTBLS2542^5^ | RP – ESI – NEG | Thermo Q Exactive Orbitrap | 5 | 161 | 299 | <https://zenodo.org/record/5804501> |
| IROA standards | GNPS | MSV000088661 | RP – ESI – POS | Thermo Fusion Orbitrap | 7.5 | 58 | 367 | <https://zenodo.org/record/5803968> |
| Cardiovascular disease | MetabolomicsWorkbench | ST002044 | RP – ESI – POS/NEG | Thermo Fusion Orbitrap | 7.5 | 286 | N/A | <https://zenodo.org/record/5803970> |
| Gut microbiota | MetabolomicsWorkbench | ST001683^6^ | RP – ESI – POS/NEG | Agilent 6545 QTOF | 10 | 210/256 | 307/273 | <https://zenodo.org/record/5804558> |
| Mouse plasma | MetabolomicsWorkbench | ST001154^7^ | HILIC – ESI – POS | Thermo Q Exactive Plus Orbitrap | 10 | 226 | 112 | <https://zenodo.org/record/5804523> |
| Pregnancy dynamics | MetabolomicsWorkbench | ST001430^8^ | RP – ESI – POS/NEG | Thermo Q Exactive Plus Orbitrap | 5 | 781 | N/A | <https://zenodo.org/record/5804527> |
| * Experimental details for LC-HRMS data collection are available in the corresponding accession ID entries (hyperlinked) in repositories. | | | | | | | | |

### S.1. Objective functions for the score coefficients optimization

1) **Overall ranking OF** may be used when number of candidate molecular formulas are high and usually true candidates cannot reach the top first candidate. Therefore, improving the overall ranking is the objective of this method.

$$OF=\min\left( \sum_{i=1}^{N_{ref}} \frac{R_{i}}{NC_{i}} \right)$$

(1)

where *R_i_*, *NC_i_* and *N_ref_* represent the rank of true compounds, number of annotated molecular formulas for the chromatographic peak and total number of reference compounds.

$$OF_{min}=\min\left( \sum_{i=1}^{N_{ref}} \frac{1}{NC_{i}} \right)$$

(2)

$$RR(\%)=\frac{OF_{unopt}-OF_{opt}}{OF_{unopt}-OF_{min}}*100\%$$

(3)

where *OF_min_*, *OF_opt_*, *OF_unopt_* and *RR* represent the minimum value of the objective function, the value of the optimized objective function, the value of the unoptimized objective function and recall rate of the optimization process, respectively.

2) **Top ranking OF** should be used when number of candidate molecular formula is significantly reduced after applying a narrow mass window. This objective function was designed to minimize the number of top candidate molecular formulas.

$$OF=\max\left( \sum_{i=1}^{N_{ref}} S(R_{i}) \right)$$

$$S(R_{i})=\left\{ \begin{aligned} 1, &R_{i}\leq maxR \\ 0, &R_{i}>maxR \end{aligned} \right.$$

(4)

where *maxR* is the maximum allowed order rank for objective function optimization. Likewise, the recall rate of the optimization process can be calculated using equation (5).

$$RR(\%)=\frac{OF_{opt}-OF_{unopt}}{OF_{max}-OF_{unopt}}*100\%$$

(5)

where *OF_max_* represents the maximum value of the objective function and simply can be measured (*OF_max_*=*N_ref_*).

The IDSL.UFA software employs the genetic algorithm R package^9^ to compute *coeff[1-5]* when a list of reference compounds are available. A specific tab in the tutorial IDSL.UFA spreadsheet is also allocated to the optimization of the score function coefficients.

| **Table S.2.** Polymeric carbon chain progression for PFAS | | | | | | |
| --- | --- | --- | --- | --- | --- | --- |
| **Structures *** | **Molecular Ion** | **C** | **∆C** ** | **F** | **∆F** ** | **∆F/∆C** |
| 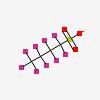 | [C_4_F_9_O_3_S]^-^ | 4 |  | 9 |  |  |
| 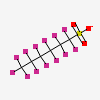 | [C_6_F_13_O_3_S]^-^ | 6 | 6 - 4 = 2 | 13 | 13 - 9 = 4 | 2 |
| 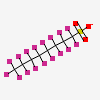 | [C_8_F_17_O_3_S]^-^ | 8 | 8 - 6 = 2 | 17 | 17 - 13 = 4 | 2 |
| 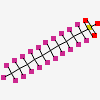 | [C_10_F_21_O_3_S]^-^ | 10 | 10 - 8 = 2 | 21 | 21 - 17 = 4 | 2 |
| 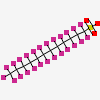 | [C_12_F_25_O_3_S]^-^ | 12 | 12 - 10 = 2 | 25 | 25 - 21 = 4 | 2 |
| * Structure images are adapted from <https://pubchem.ncbi.nlm.nih.gov/>  ** **∆C** and **∆F** represent difference in number of carbon and fluorine atoms in nearby rows. | | | | | | |

| **Table S.3.** Cyclic chain progression for polycyclic phenols | | | | | | |
| --- | --- | --- | --- | --- | --- | --- |
| **Structures *** | **Molecular formula** | **C** | **∆C** ** | **H** | **∆H** ** | **∆H/∆C** |
| 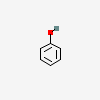 | C_6_H_6_O | 6 |  | 6 |  |  |
| 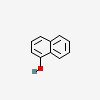 | C_10_H_8_O | 10 | 10 - 6 = 4 | 8 | 8 - 6 = 2 | 1/2 |
| 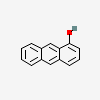 | C_14_H_10_O | 14 | 14 - 10 = 4 | 10 | 10 - 8 = 2 | 1/2 |
| 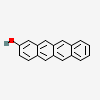 | C_18_H_12_O | 18 | 18 - 14 = 4 | 12 | 12 - 10 = 2 | 1/2 |
| 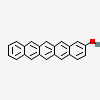 | C_22_H_14_O | 22 | 22 - 18 = 4 | 14 | 14 - 12 = 2 | 1/2 |
| * Structure images were adapted from <https://pubchem.ncbi.nlm.nih.gov/>  ** **∆C** and **∆H** represent difference in number of carbon and fluorine atoms in nearby rows. | | | | | | |

| **Table S.4.** Repeated linear and cyclic groups in a class of compounds | | | | | |
| --- | --- | --- | --- | --- | --- |
|  |  | $\vec{\boldsymbol{linear carbon chain progress}}$ | | | |
|  | **Structures *** | 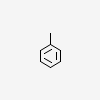 | 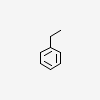 | 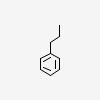 | $R=\frac{\Delta H}{\Delta C}=\frac{12-10}{9-8}=\frac{10-8}{8-7}=2$ |
| $\overset{⃖}{\boldsymbol{cyclic chain progress}}$ | 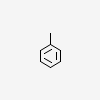 | C_7_H_8_ | C_8_H_10_ | C_9_H_12_ |  |
|  | 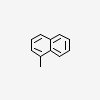 | C_11_H_10_ |  |  |  |
|  | 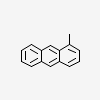 | C_15_H_12_ |  |  |  |
|  | $R=\frac{\Delta H}{\Delta C}=\frac{12-10}{15-11}=\frac{10-8}{11-7}=\frac{1}{2}$ | |  |  |  |
| * Structure images were adapted from <https://pubchem.ncbi.nlm.nih.gov/> | | | | | |

| **Table S.5.** Constant numbers of carbons and Σ(H+Cl) for PCBs | | | | | |
| --- | --- | --- | --- | --- | --- |
| **Structures*** | **Molecular Ion** | **C** | **H** | **Cl** | **Σ(H+Cl)** |
| 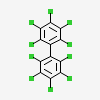 | [C_12_Cl_10_]^+^ | 12 | 0 | 10 | 0 + 10 = 10 |
| 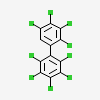 | [C_12_HCl_9_]^+^ | 12 | 1 | 9 | 1 + 9 = 10 |
| 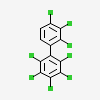 | [C_12_H_2_Cl_8_]^+^ | 12 | 2 | 8 | 2 + 8 = 10 |
| 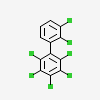 | [C_12_H_3_Cl_7_]^+^ | 12 | 3 | 7 | 3 + 7 = 10 |
| 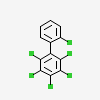 | [C_12_H_4_Cl_6_]^+^ | 12 | 4 | 6 | 4 + 6 = 10 |
| 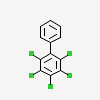 | [C_12_H_5_Cl_5_]^+^ | 12 | 5 | 5 | 5 + 5 = 10 |
| 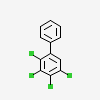 | [C_12_H_6_Cl_4_]^+^ | 12 | 6 | 4 | 6 + 4 =10 |
| 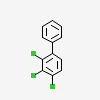 | [C_12_H_7_Cl_3_]^+^ | 12 | 7 | 3 | 7 + 3 = 10 |
| 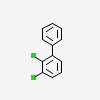 | [C_12_H_8_Cl_2_]^+^ | 12 | 8 | 2 | 8 + 2 = 10 |
| 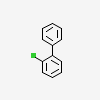 | [C_12_H_9_Cl]^+^ | 12 | 9 | 1 | 9 + 1 = 10 |
| 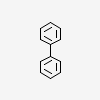 | [C_12_H_10_]^+^ | 12 | 10 | 0 | 10 + 0 = 10 |
| * Structure images were adapted from <https://pubchem.ncbi.nlm.nih.gov/> | | | | | |

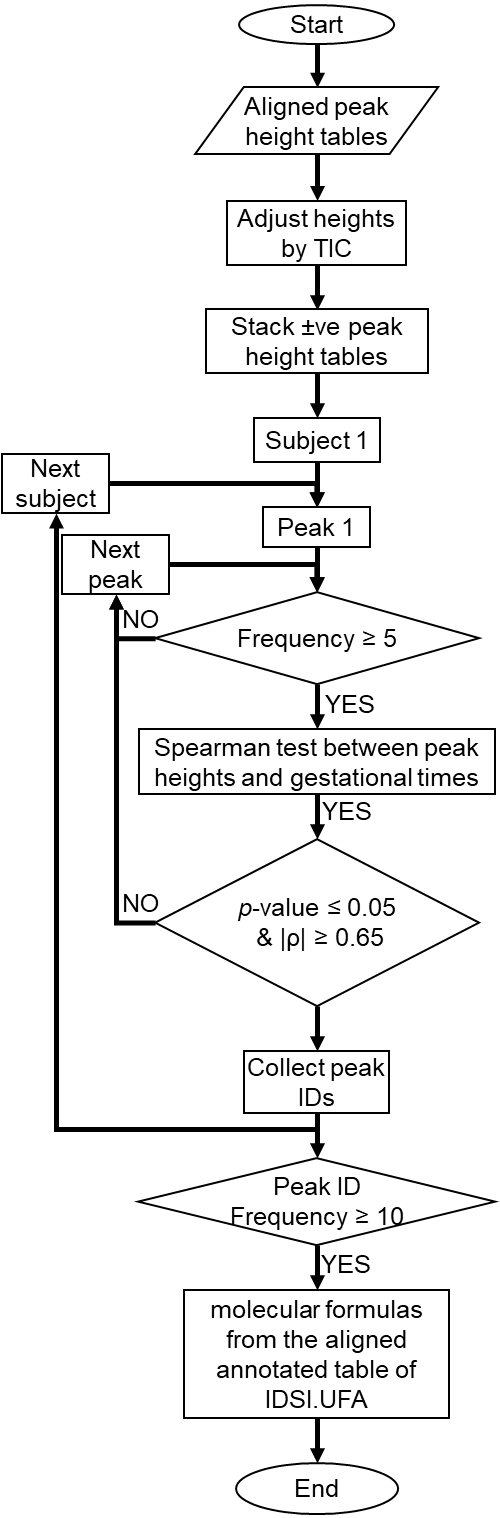

**Figure S.2.** Schematic of workflow to annotate peaks with linear patterns in pregnancy.

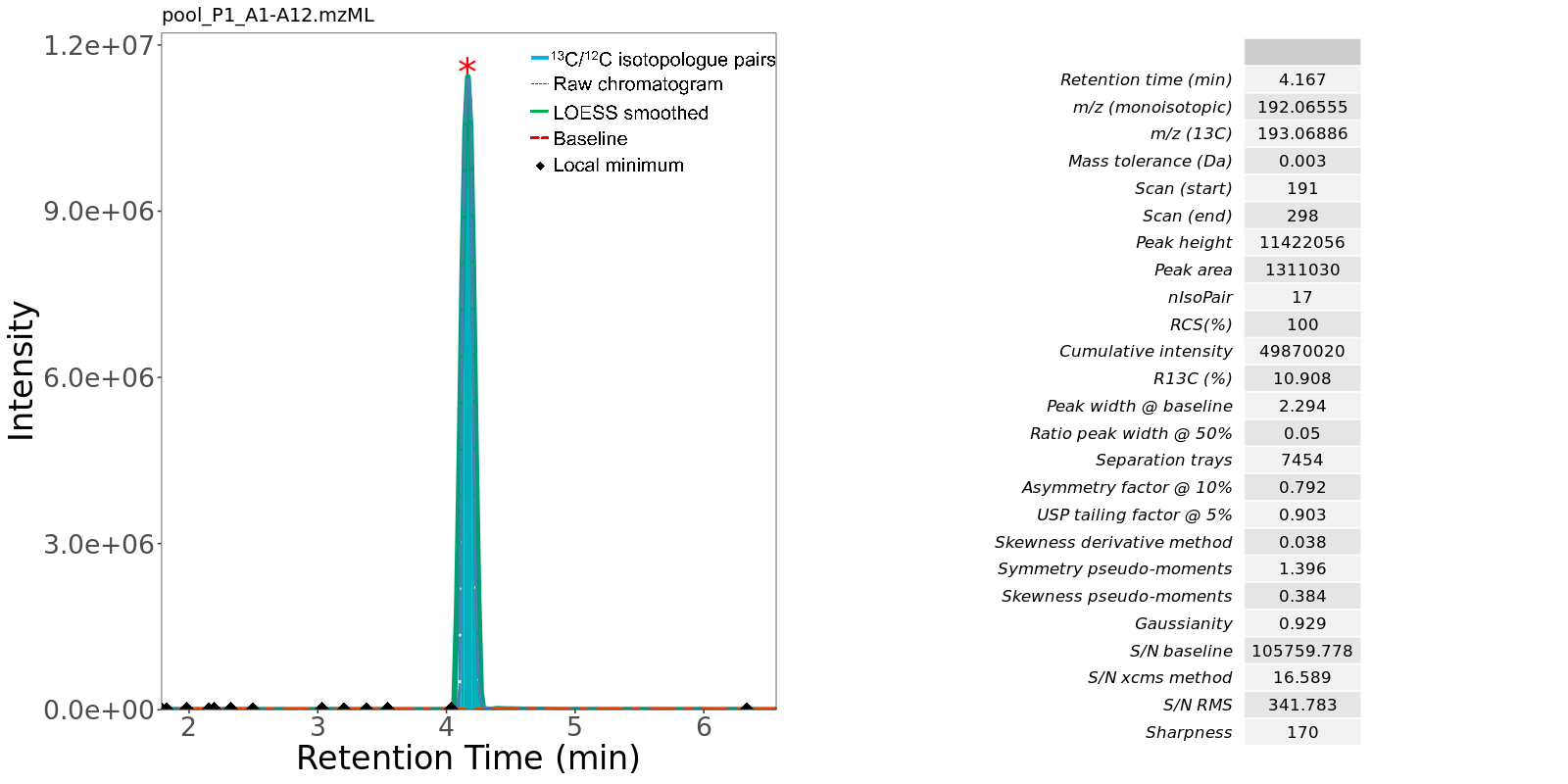

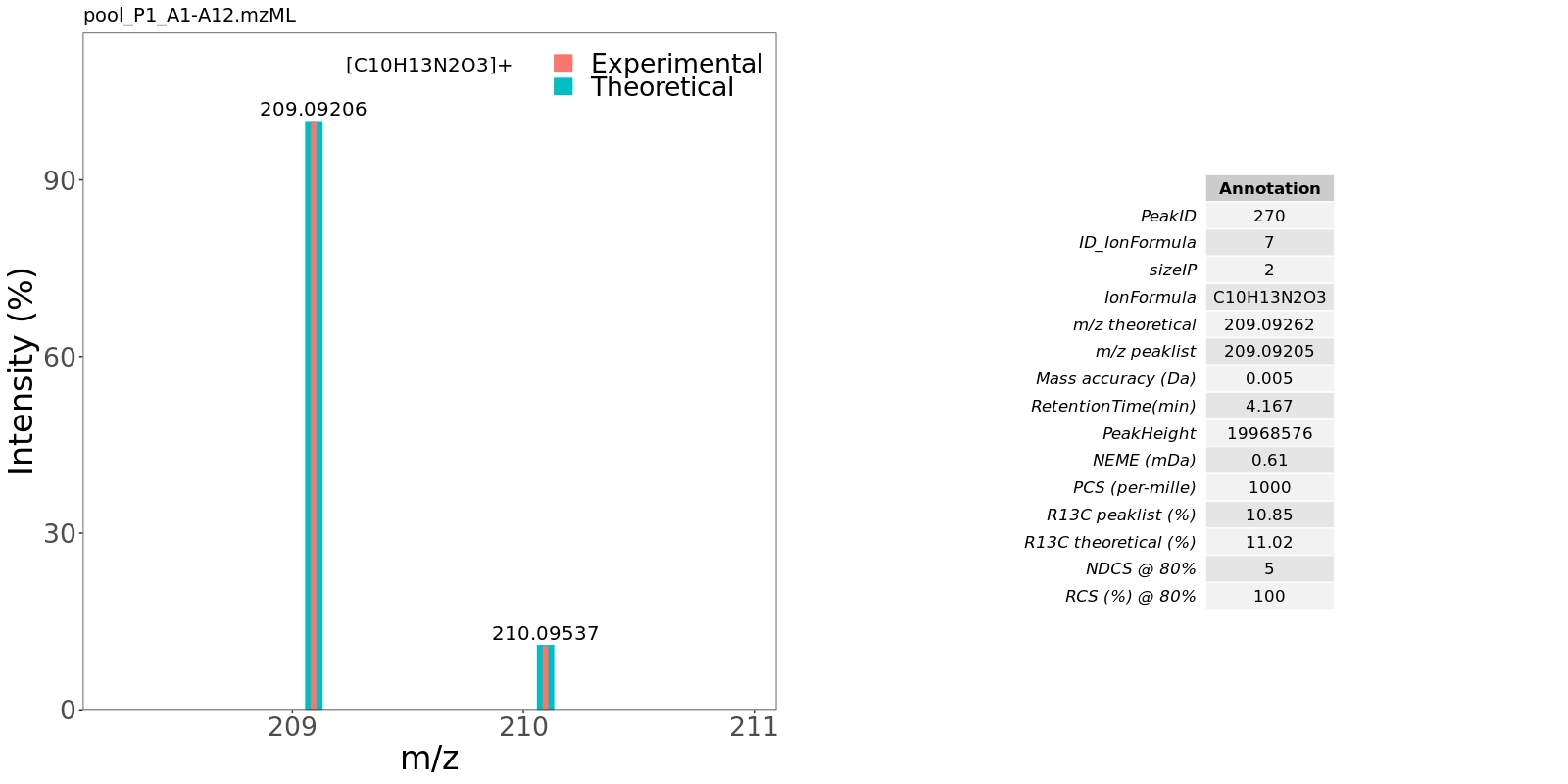

**Figure S.3.** Comparison between theoretical and integrated experimental isotopic profiles across a chromatographic peak after molecular formula annotation using the IDSL.UFA pipeline on an authentic standard from the MSV000088661 study.

| **IDSL.ExposomeDB IPDB** | **Enumerated Chemical Space (ECS) IPDBs*** |
| --- | --- |
| 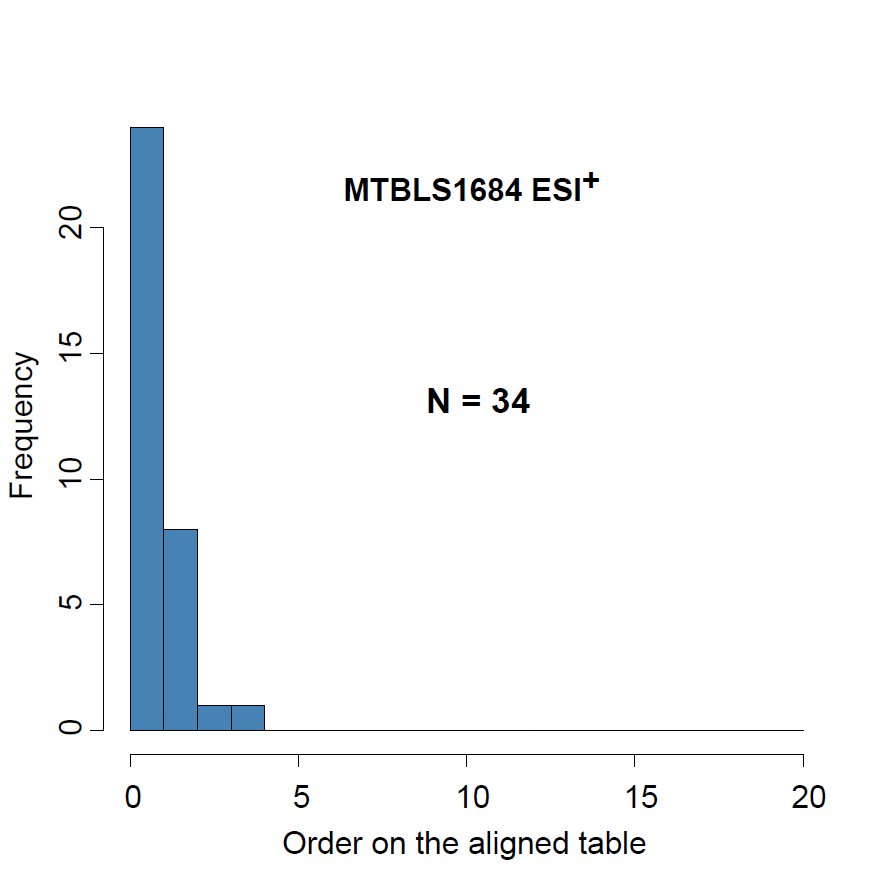 | 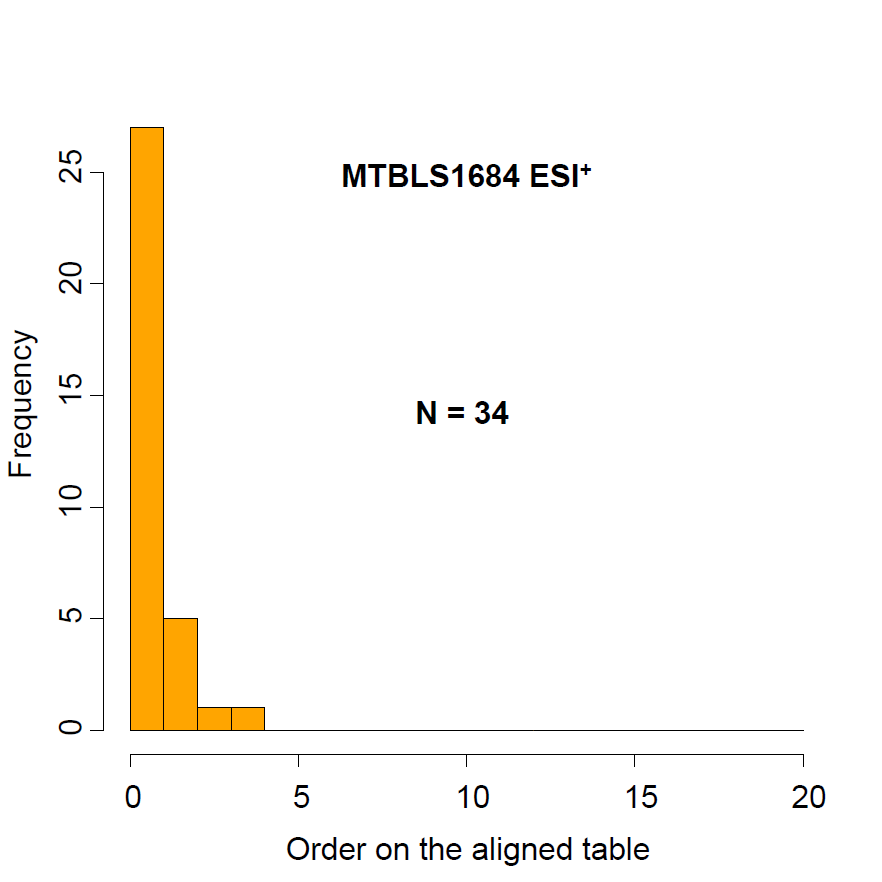 |
| 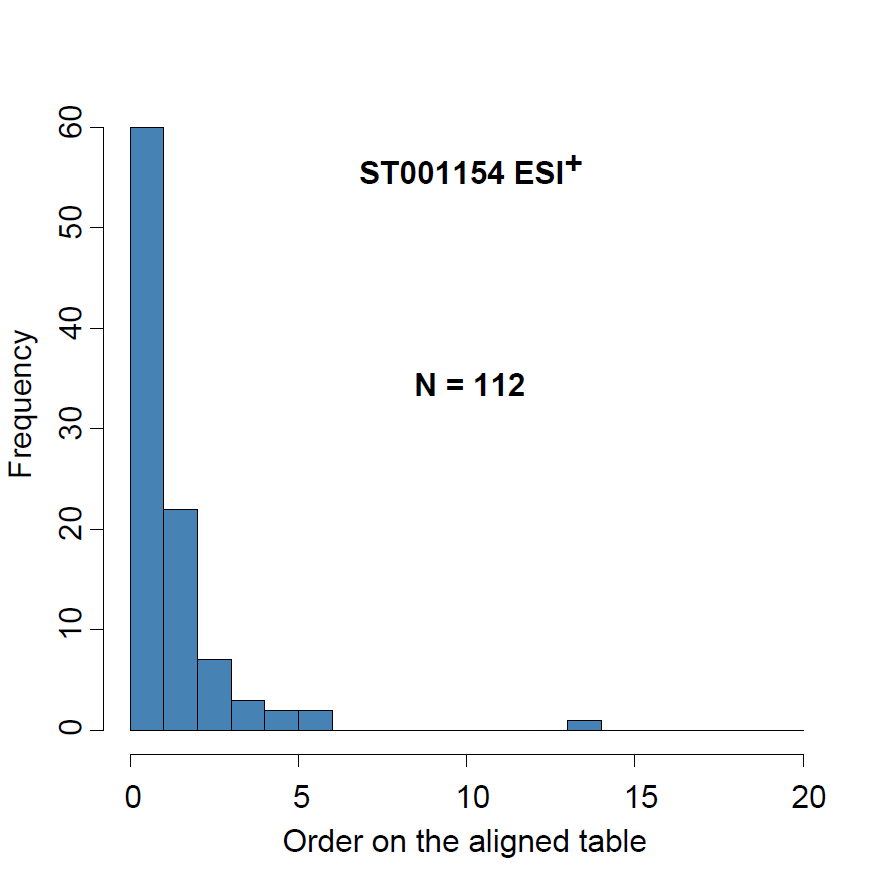 | 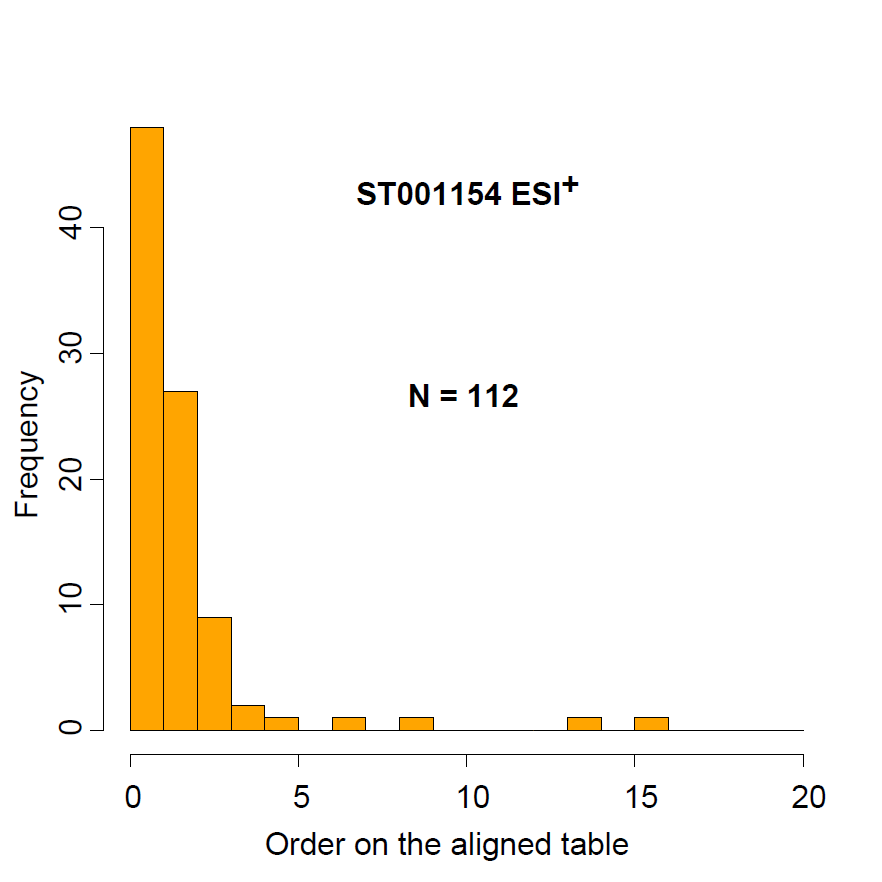 |
| 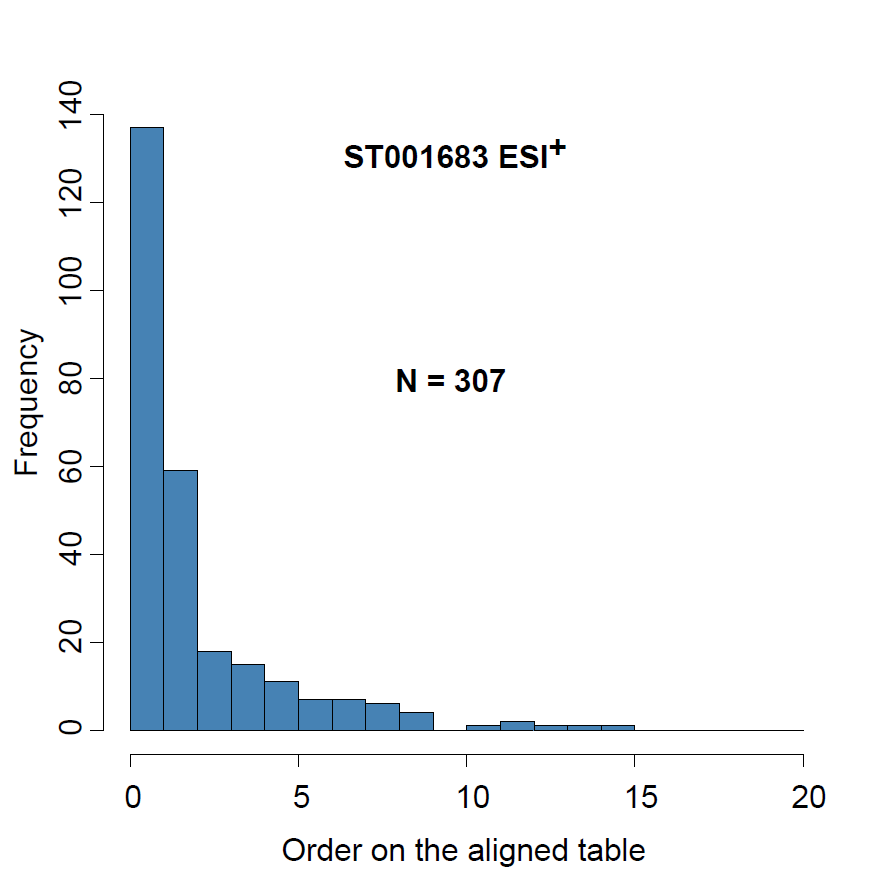 | 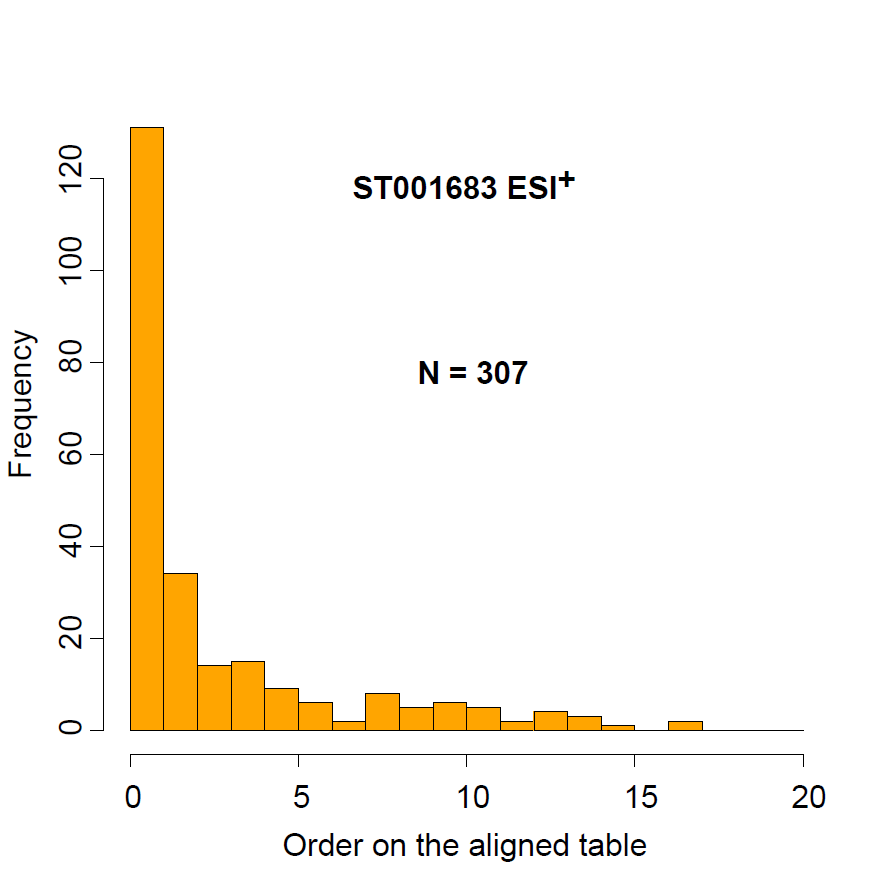 |
| 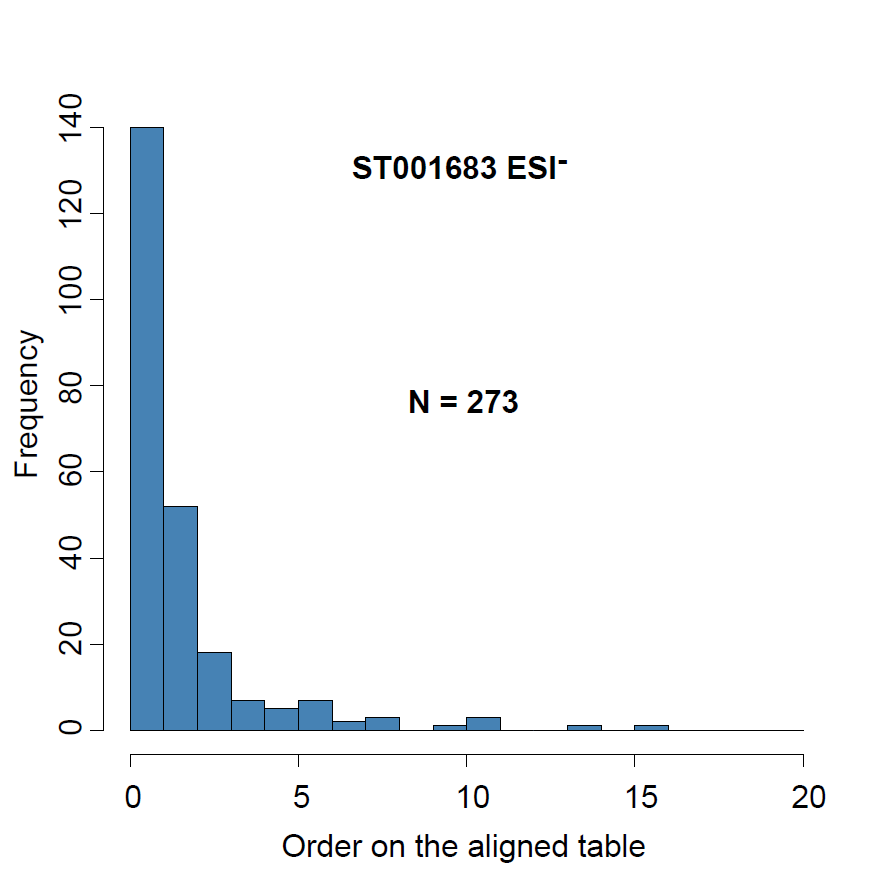 | 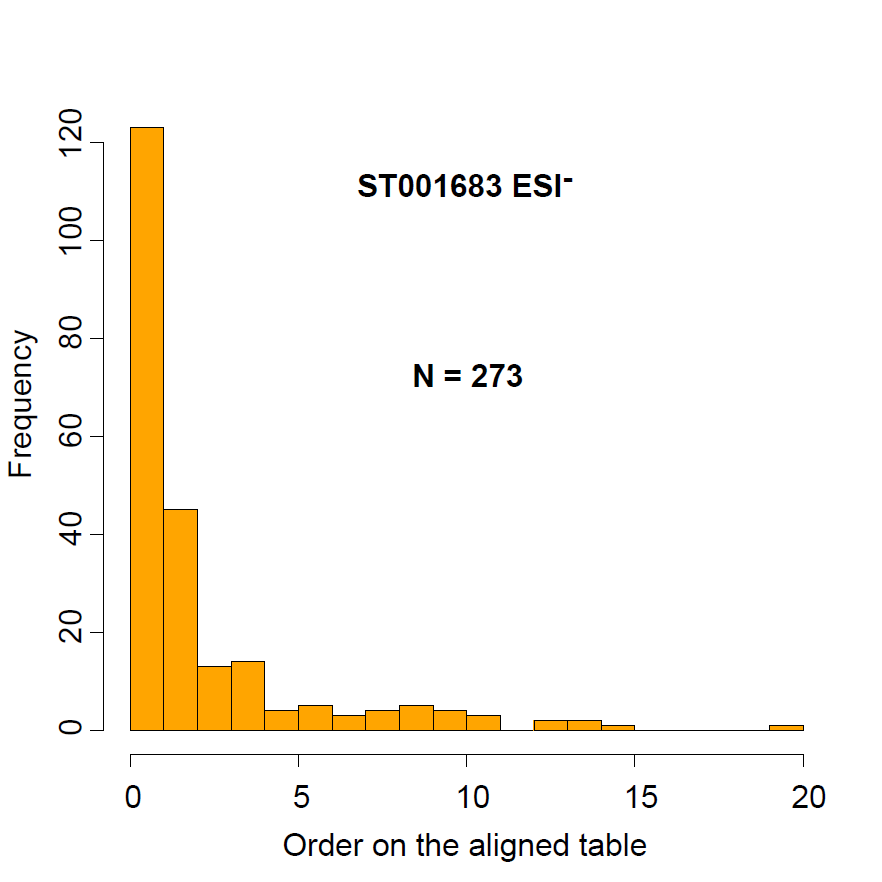 |
| 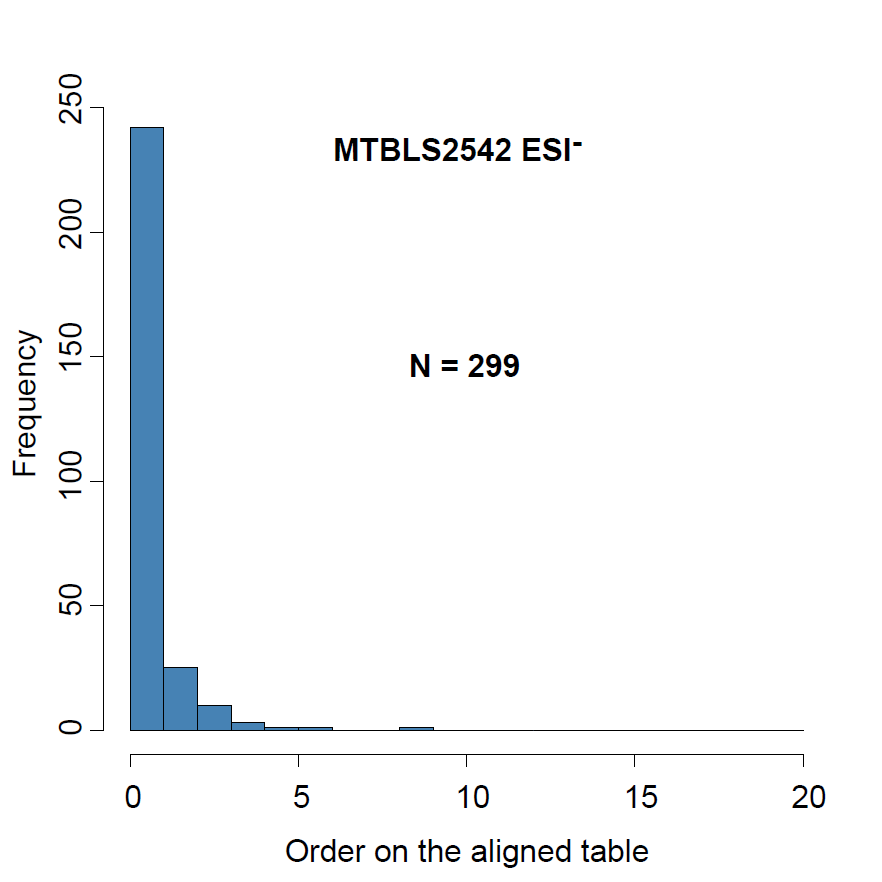 | 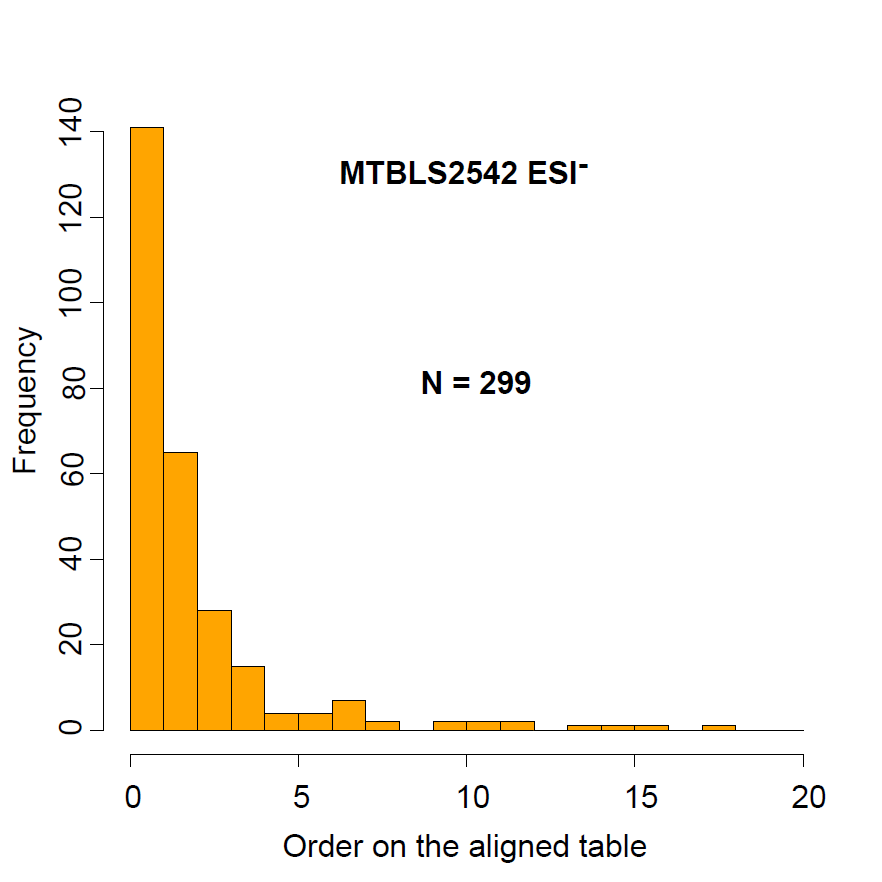 |
| **Figure S.4.** Distribution of the candidate order on the aligned annotated molecular formula tables in each study. (* IPDBs may be found at <https://zenodo.org/record/5823455>) | |

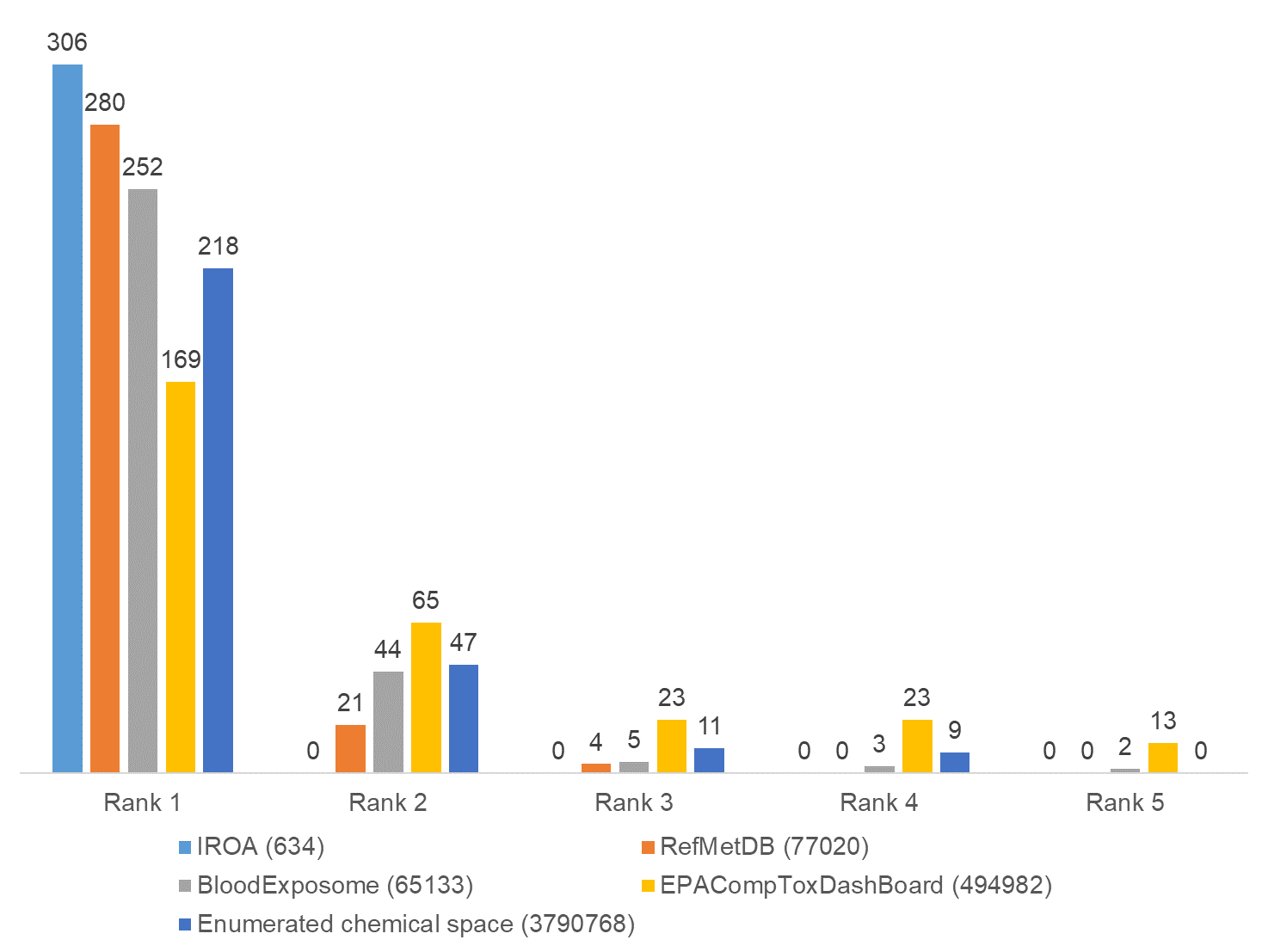

**Figure S.5.** Evaluation of performance of the ranking method for true positives (only [M+H]^+^ ionization used) with respect to various chemical spaces on the MSV000088661 study (size of chemical space is shown inside parenthesis). (https://zenodo.org/record/6609406)

| **Table S.6.** Comparison between aligned annotated table using IDSL.ExposomeDB IPDB by the IDSL.UFA and MS/MS library match for the ST002044 study. | | | | | | | | |
| --- | --- | --- | --- | --- | --- | --- | --- | --- |
| **IDSL.UFA** | | | | | | **MS/MS library match** | | |
| **Order on the aligned table** | **Median rank** | **m/z** | **ID** | **Ion formula** | **RT (min)** | **ID** | **Ion formula** | **RT (min)** |
| 1 | 1 | 167.02052 | 15186 | [C_5_H_3_N_4_O_3_]^-^ | 1.54 | 1 | [C_5_H_3_N_4_O_3_]^-^ | 1.53 |
| 3 | 3 | 568.36144 | 162747 | [C_27_H_55_NO_9_P]^-^ | 9.97 | 2 | [C_27_H_55_NO_9_P]^-^ | 9.93 |
| 1 | 1 | 564.33014 | 161867 | [C_27_H_51_NO_9_P]^-^ | 10.87 | 3 | [C_27_H_51_NO_9_P]^-^ | 10.97 |
| 2 | 2 | 802.55981 | 190802 | [C_43_H_81_NO_10_P]^-^ | 10.81 | 4 | [C_43_H_81_NO_10_P]^-^ | 10.88 |
| 1 | 1 | 311.13957 | 76831 | [C_18_H_19_N_2_O_3_]^-^ | 7.77 | 5 | [C_18_H_19_N_2_O_3_]^-^ | 7.79 |
| 1 | 1 | 204.06607 | 31026 | [C_11_H_10_NO_3_]^-^ | 7.78 | 6 | [C_11_H_10_NO_3_]^-^ | 7.79 |
| 2 | 2 | 554.34579 | 159698 | [C_26_H_53_NO_9_P]^-^ | 11.46 | 7 | [C_26_H_53_NO_9_P]^-^ | 11.47 |
| 2 | 2 | 804.57546 | 190948 | [C_43_H_83_NO_10_P]^-^ | 10.99 | 8 | [C_43_H_83_NO_10_P]^-^ | 11.01 |
| 1 | 1 | 307.02763 | 74829 | [C_14_H_11_O_6_S]^-^ | 8.24 | 9 | [C_14_H_11_O_6_S]^-^ | 8.25 |
| 1 | 1 | 191.01918 | 25546 | [C_6_H_7_O_7_]^-^ | 1.31 | 10 | [C_6_H_7_O_7_]^-^ | 1.31 |
| 2 | 2 | 382.06731 | 105326 | [C_16_H_14_F_2_N_3_O_4_S]^-^ | 8.54 | 11 | [C_16_H_14_F_2_N_3_O_4_S]^-^ | 8.57 |
| 1 | 1 | 197.01985 | 27732 | [C_7_H_5_N_2_O_5_]^-^ | 9.11 | 12 | [C_7_H_5_N_2_O_5_]^-^ | 9.14 |
| 1 | 1 | 164.07115 | 14276 | [C_9_H_10_NO_2_]^-^ | 4.64 | 13 | [C_9_H_10_NO_2_]^-^ | 4.63 |
| 2 | 2 | 279.09810 | 62565 | [C_13_H_15_N_2_O_5_]^-^ | 5.87 | 14 | [C_13_H_15_N_2_O_5_]^-^ | 5.89 |
| 1 | 1 | 410.08221 | 115444 | [C_17_H_17_FN_3_O_6_S]^-^ | 8.15 | 15 | [C_17_H_17_FN_3_O_6_S]^-^ | 8.18 |
| 1 | 2 | 326.08759 | 83074 | [C_14_H_16_NO_8_]^-^ | 4.35 | 16 | [C_14_H_16_NO_8_]^-^ | 4.33 |
| 1 | 1 | 478.29336 | 138111 | [C_23_H_45_NO_7_P]^-^ | 11.29 | 17 | [C_23_H_45_NO_7_P]^-^ | 11.30 |
| 1 | 1 | 325.18374 | 82767 | [C_18_H_29_O_3_S]^-^ | 10.77 | 18 | [C_18_H_29_O_3_S]^-^ | 10.96 |
| 3 | 3 | 432.20223 | 122847 | [C_23_H_30_NO_7_]^-^ | 5.8 | 19 | [C_23_H_30_NO_7_]^-^ | 5.84 |
| 2 | 2 | 195.05182 | 26828 | [C_7_H_7_N_4_O_3_]^-^ | 5.75 | 20 | [C_7_H_7_N_4_O_3_]^-^ | 5.76 |
| 2 | 2 | 364.09672 | 98437 | [C_16_H_18_N_3_O_5_S]^-^ | 4.98 | 21 | [C_16_H_18_N_3_O_5_S]^-^ | 5.01 |
| 1 | 1 | 180.06607 | 21197 | [C_9_H_10_NO_3_]^-^ | 2.36 | 22 | [C_9_H_10_NO_3_]^-^ | 2.39 |
| 2 | 2 | 512.29884 | 148571 | [C_23_H_47_NO_9_P]^-^ | 10.52 | 23 | [C_23_H_47_NO_9_P]^-^ | 10.53 |
| 1 | 1 | 319.22732 | 80409 | [C_20_H_31_O_3_]^-^ | 10.54 | 24 | [C_20_H_31_O_3_]^-^ | 10.56 |
| 1 | 1 | 407.27975 | 114675 | [C_24_H_39_O_5_]^-^ | 10.16 | 25 | [C_24_H_39_O_5_]^-^ | 10.17 |
| 3 | 3 | 405.26410 | 114021 | [C_24_H_37_O_5_]^-^ | 9.66 | 26 | [C_24_H_37_O_5_]^-^ | 9.69 |
| 2 | 2 | 481.28014 | 138972 | [C_26_H_41_O_8_]^-^ | 10.03 | 27 | [C_26_H_41_O_8_]^-^ | 10.06 |
| 1 | 1 | 595.28834 | 168381 | [C_27_H_48_O_12_P]^-^ | 11.46 | 28 | [C_27_H_48_O_12_P]^-^ | 11.46 |
| 4 | 4 | 826.55981 | 192714 | [C_45_H_81_NO_10_P]^-^ | 10.46 | 29 | [C_45_H_81_NO_10_P]^-^ | 10.54 |
| 1 | 1 | 309.10866 | 75961 | [C_14_H_17_N_2_O_6_]^-^ | 4.49 | 30 | [C_14_H_17_N_2_O_6_]^-^ | 4.99 |
| 2 | 2 | 309.10866 | 75960 | [C_14_H_17_N_2_O_6_]^-^ | 4.99 | 31 | [C_14_H_17_N_2_O_6_]^-^ | 5.00 |
| 2 | 2 | 194.04533 | 26418 | [C_9_H_8_NO_4_]^-^ | 5.13 | 32 | [C_9_H_8_NO_4_]^-^ | 5.15 |
| 1 | 1 | 221.09262 | 37312 | [C_11_H_13_N_2_O_3_]^-^ | 5.67 | 33 | [C_11_H_13_N_2_O_3_]^-^ | 5.65 |
| 2 | 2 | 251.10318 | 50699 | [C_12_H_15_N_2_O_4_]^-^ | 4.35 | 34 | [C_12_H_15_N_2_O_4_]^-^ | 4.35 |
| 4 | 3.5 | 445.19881 | 127307 | [C_24_H_25_N_6_O_3_]^-^ | 8.28 | 35 | [C_24_H_25_N_6_O_3_]^-^ | 8.29 |
| 1 | 1 | 310.00377 | 76395 | [C_14_H_10_Cl_2_NO_3_]^-^ | 9.31 | 36 | [C_14_H_10_Cl_2_NO_3_]^-^ | 9.31 |
| 1 | 1 | 259.12940 | 54112 | [C_11_H_19_N_2_O_5_]^-^ | 6.24 | 37 | [C_11_H_19_N_2_O_5_]^-^ | 6.24 |
| 2 | 2 | 436.28280 | 124105 | [C_21_H_43_NO_6_P]^-^ | 11.45 | 38 | [C_21_H_43_NO_6_P]^-^ | 11.47 |
| 1 | 1 | 557.24518 | 160414 | [C_33_H_34_FN_2_O_5_]^-^ | 9.56 | 39 | [C_33_H_34_FN_2_O_5_]^-^ | 9.60 |
| 6 | 5.5 | 323.13957 | 81910 | [C_19_H_19_N_2_O_3_]^-^ | 9.17 | 40 | [C_19_H_19_N_2_O_3_]^-^ | 9.15 |
| 1 | 1 | 307.14465 | 75021 | [C_19_H_19_N_2_O_2_]^-^ | 9.66 | 41 | [C_19_H_19_N_2_O_2_]^-^ | 9.68 |
| 1 | 1 | 340.14096 | 88668 | [C_17_H_18_N_5_O_3_]^-^ | 4.84 | 42 | [C_17_H_18_N_5_O_3_]^-^ | 4.85 |
| 1 | 1 | 291.06573 | 67868 | [C_18_H_11_O_4_]^-^ | 9.39 | 43 | [C_18_H_11_O_4_]^-^ | 9.39 |
| 2 | 2 | 256.09737 | 52674 | [C_15_H_14_NO_3_]^-^ | 9.35 | 44 | [C_15_H_14_NO_3_]^-^ | 9.36 |
| 1 | 1 | 337.00498 | 87250 | [C_14_H_10_ClN_2_O_4_S]^-^ | 7.8 | 45 | [C_14_H_10_ClN_2_O_4_S]^-^ | 7.84 |
| 1 | 1 | 227.10318 | 39921 | [C_10_H_15_N_2_O_4_]^-^ | 5.71 | 46 | [C_10_H_15_N_2_O_4_]^-^ | 5.72 |
| 1 | 1 | 179.05690 | 20937 | [C_7_H_7_N_4_O_2_]^-^ | 6 | 47 | [C_7_H_7_N_4_O_2_]^-^ | 6.00 |
| 4 | 4 | 830.59111 | 193014 | [C_45_H_85_NO_10_P]^-^ | 11.3 | 48 | [C_45_H_85_NO_10_P]^-^ | 11.32 |
| 1 | 1 | 277.15522 | 61740 | [C_15_H_21_N_2_O_3_]^-^ | 7.62 | 49 | [C_15_H_21_N_2_O_3_]^-^ | 7.62 |
| 3 | 2.5 | 480.16046 | 138595 | [C_22_H_27_FN_3_O_6_S]^-^ | 9.11 | 50 | [C_22_H_27_FN_3_O_6_S]^-^ | 9.13 |
| 1 | 1 | 347.11779 | 91498 | [C_16_H_19_N_4_O_3_S]^-^ | 8.5 | 51 | [C_16_H_19_N_4_O_3_S]^-^ | 8.51 |
| 1 | 1 | 179.07082 | 20969 | [C_10_H_11_O_3_]^-^ | 9.15 | 52 | [C_10_H_11_O_3_]^-^ | 9.16 |
| 2 | 2 | 495.29579 | 143129 | [C_27_H_43_O_8_]^-^ | 9.95 | 53 | [C_27_H_43_O_8_]^-^ | 9.97 |
| 1 | 1 | 197.06747 | 27908 | [C_7_H_9_N_4_O_3_]^-^ | 1.22 | 54 | [C_7_H_9_N_4_O_3_]^-^ | 1.25 |
| 2 | 2 | 566.34579 | 162295 | [C_27_H_53_NO_9_P]^-^ | 11.67 | 55 | [C_27_H_53_NO_9_P]^-^ | 11.52 |
| 2 | 2 | 465.24884 | 133990 | [C_25_H_37_O_8_]^-^ | 9.66 | 56 | [C_25_H_37_O_8_]^-^ | 9.67 |
| 1 | 1 | 261.00690 | 54836 | [C_9_H_9_O_7_S]^-^ | 5.44 | 57 | [C_9_H_9_O_7_S]^-^ | 5.47 |
| 2 | 2 | 288.14471 | 66695 | [C_13_H_22_NO_6_]^-^ | 4.85 | 58 | [C_13_H_22_NO_6_]^-^ | 4.84 |
| 1 | 2 | 181.03617 | 21479 | [C_6_H_5_N_4_O_3_]^-^ | 4.44 | 59 | [C_6_H_5_N_4_O_3_]^-^ | 4.45 |
| 9 | 9 | 513.22905 | 148766 | [C_33_H_29_N_4_O_2_]^-^ | 9.14 | 60 | [C_33_H_29_N_4_O_2_]^-^ | 9.12 |
| 1 | 1 | 330.05485 | 84780 | [C_15_H_12_N_3_O_4_S]^-^ | 8.98 | 61 | [C_15_H_12_N_3_O_4_S]^-^ | 8.98 |
| 1 | 1 | 461.07200 | 132650 | [C_21_H_17_O_12_]^-^ | 8.33 | 62 | [C_21_H_17_O_12_]^-^ | 8.35 |
| 1 | 1 | 434.21921 | 123533 | [C_24_H_28_N_5_O_3_]^-^ | 9.46 | 63 | [C_24_H_28_N_5_O_3_]^-^ | 9.47 |
| 1 | 1 | 450.21413 | 128979 | [C_24_H_28_N_5_O_4_]^-^ | 8.88 | 64 | [C_24_H_28_N_5_O_4_]^-^ | 8.90 |
| 1 | 1 | 302.11408 | 72785 | [C_15_H_16_N_3_O_4_]^-^ | 6.86 | 65 | [C_15_H_16_N_3_O_4_]^-^ | 6.86 |
| 1 | 1 | 156.99594 | 11706 | [C_6_H_5_O_3_S]^-^ | 3.48 | 66 | [C_6_H_5_O_3_S]^-^ | 3.47 |
| 2 | 2 | 221.08138 | 37255 | [C_12_H_13_O_4_]^-^ | 8.68 | 67 | [C_12_H_13_O_4_]^-^ | 9.17 |
| 1 | 1 | 221.08138 | 37296 | [C_12_H_13_O_4_]^-^ | 9.21 | 68 | [C_12_H_13_O_4_]^-^ | 9.20 |
| 1 | 1 | 183.02935 | 22192 | [C_8_H_7_O_5_]^-^ | 5.94 | 69 | [C_8_H_7_O_5_]^-^ | 5.96 |
| 9 | 8 | 431.20698 | 122536 | [C_24_H_31_O_7_]^-^ | 8.43 | 70 | [C_24_H_31_O_7_]^-^ | 8.45 |
| 1 | 1 | 207.07697 | 31950 | [C_10_H_11_N_2_O_3_]^-^ | 4.34 | 71 | [C_10_H_11_N_2_O_3_]^-^ | 4.36 |
| 1 | 1 | 368.28008 | 100014 | [C_21_H_38_NO_4_]^-^ | 11.35 | 72 | [C_21_H_38_NO_4_]^-^ | 11.38 |
| 1 | 2 | 315.07161 | 78417 | [C_13_H_15_O_9_]^-^ | 4.93 | 73 | [C_13_H_15_O_9_]^-^ | 4.95 |

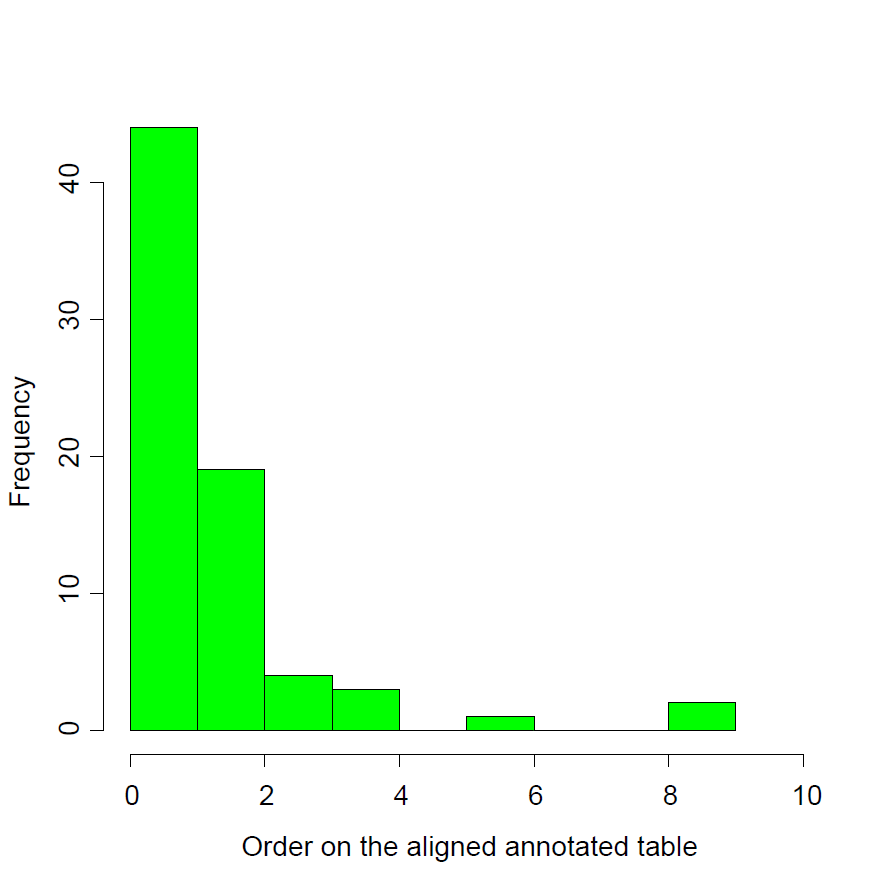
**Figure S.6.** Distribution of the candidate order on the aligned annotated molecular formula table using IDSL.ExposomeDB IPDB for the matched compounds using the MS/MS library shown on Table S.7 for the ST002044 study. A set of relaxed criteria (generally mass accuracy ≤ 7.5 mDa, $\bar{NEME}$ ≤ 7.5 mDa, $\bar{PCS}$ ≥ 95%) was used for this study.

| **Table S.7.** Evaluation of performances of the objective functions on different datasets | | | | | | | | | |
| --- | --- | --- | --- | --- | --- | --- | --- | --- | --- |
| **Study** | **Number of reference molecular formulas*** | **Objective function type** | **RR (%)** | **Number of compounds in each rank range** | | | | | |
|  |  |  |  | **Rank ≤ 1** | | **Rank ≤ 5** | | **Rank ≤ 10** | |
|  |  |  |  | **unoptimized** | **optimized** | **unoptimized** | **optimized** | **unoptimized** | **optimized** |
| MSV000088661 | 367 | Top Rank ≤ 1 | 48.48 | 268 | 316 | 364 | 364 | 366 | 365 |
| ST002044 | 3170 | Overall Rank | 48.79 | 908 | 1205 | 1608 | 2521 | 2780 | 3119 |
| * Molecular formulas may be repeated in some samples | | | | | | | | | |

| 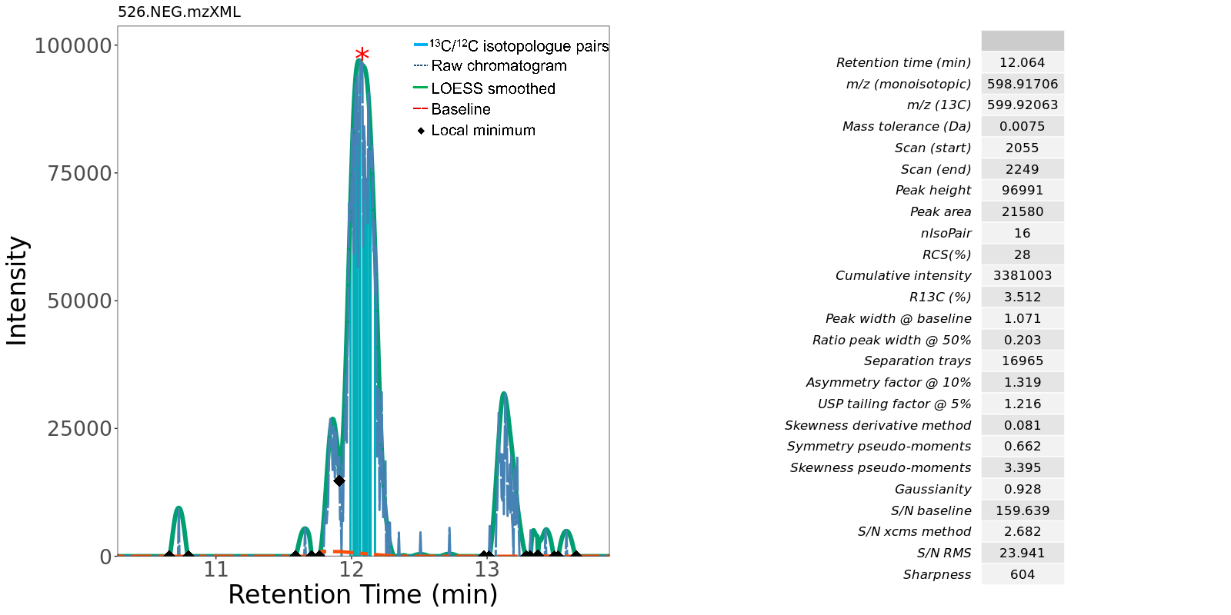 | 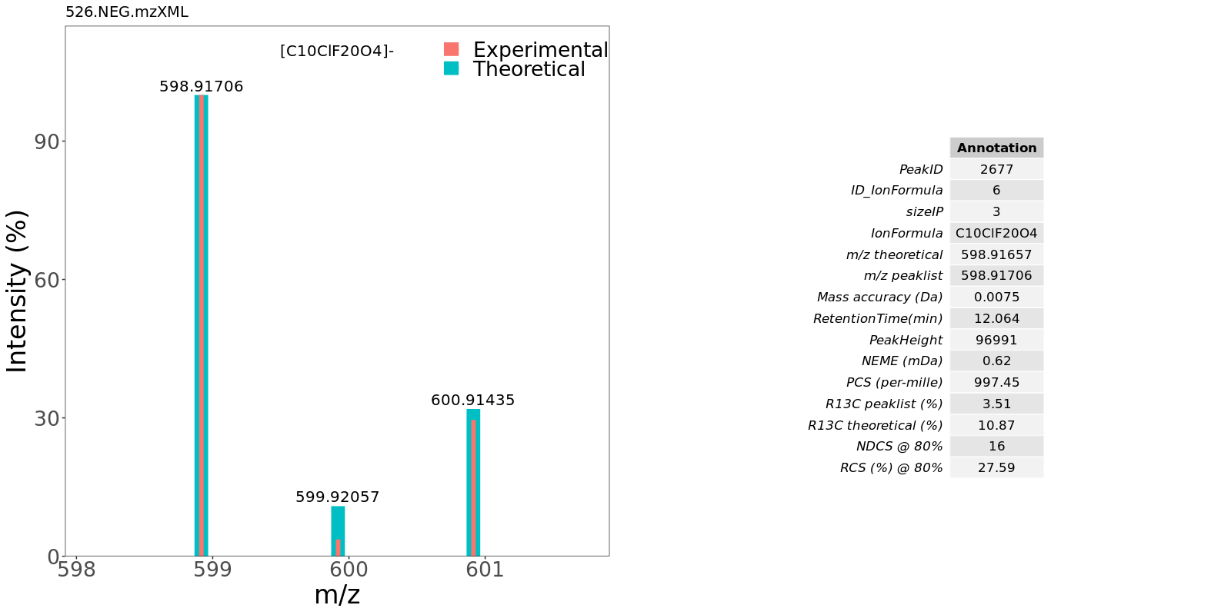 |
| --- | --- |
| 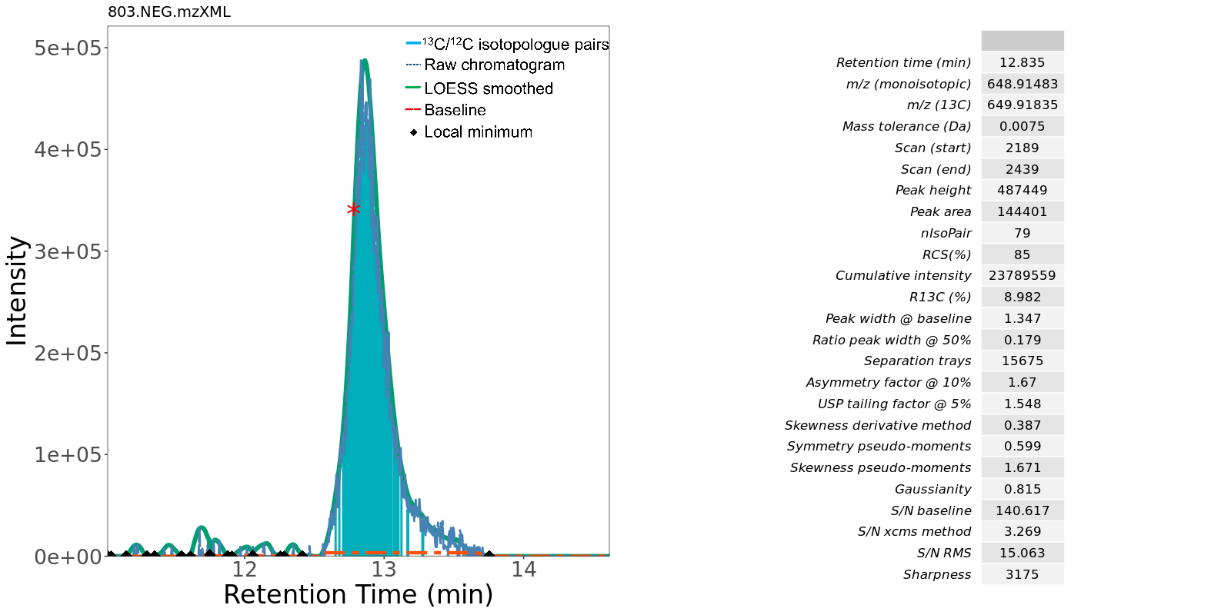 | 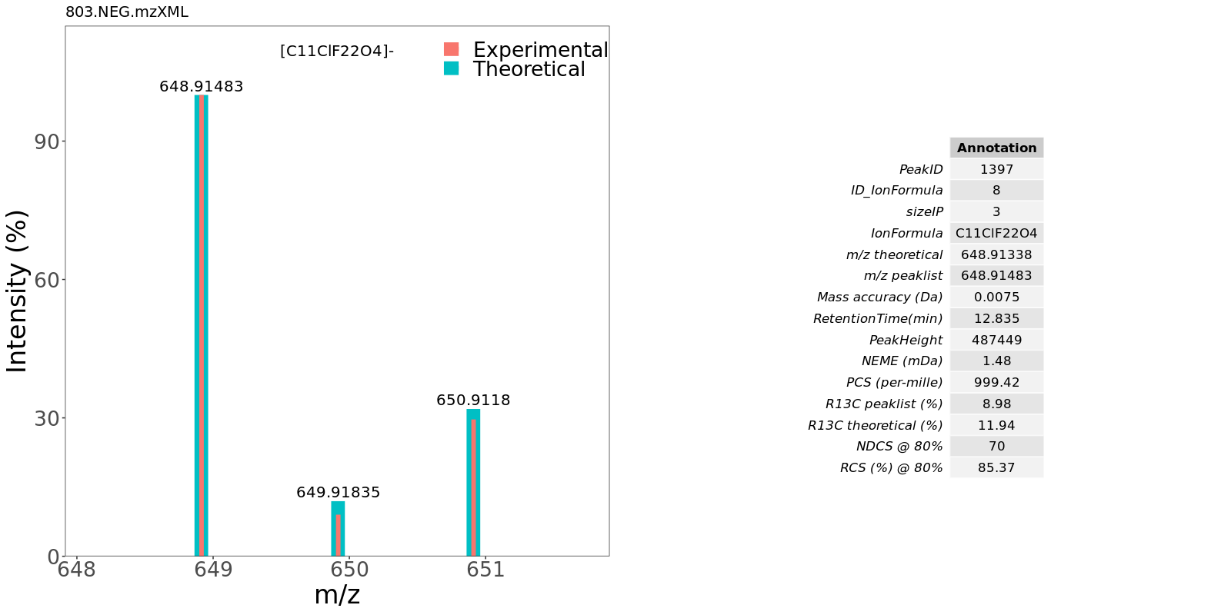 |
| 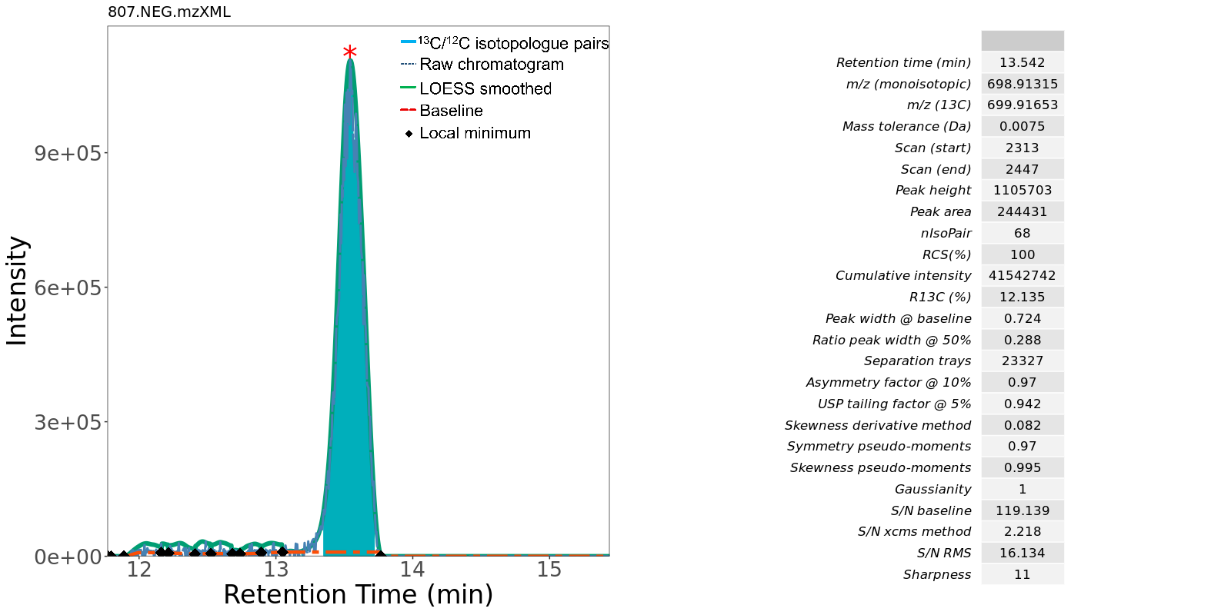 | 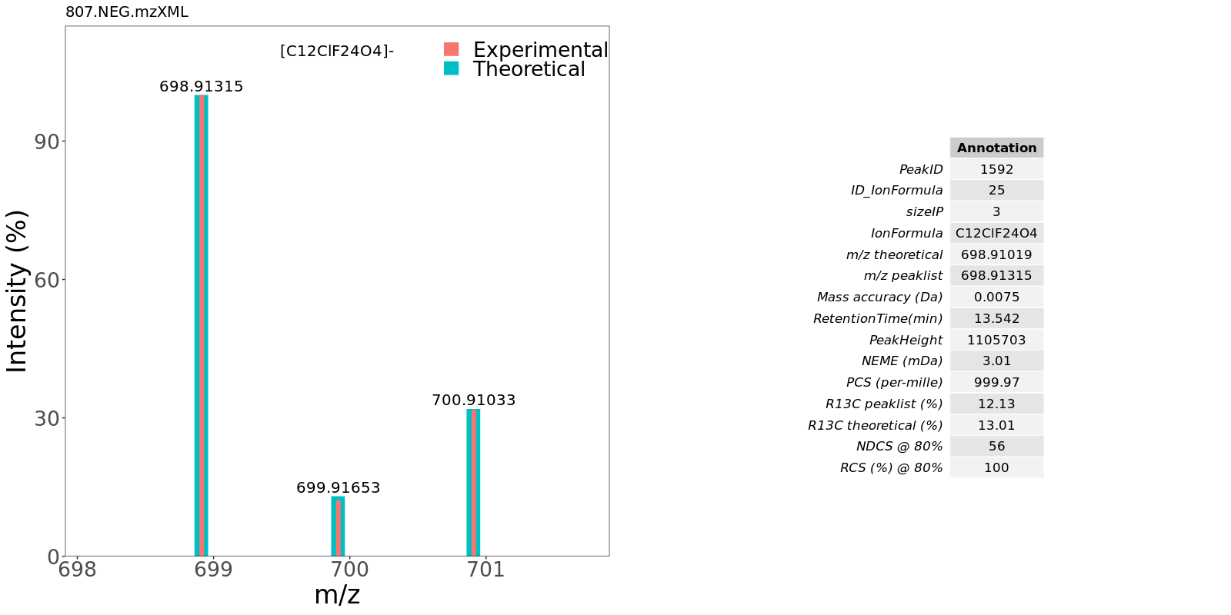 |
| **Figure S.7.** Examples of the chromatographic peaks and integrated mass spectra pertaining to the Cl-PFTrEAs ([C*_n_*ClF_2_*_n_*O_4_]^-^ (where *n* = 10-12) class in the ST001430 study. | |

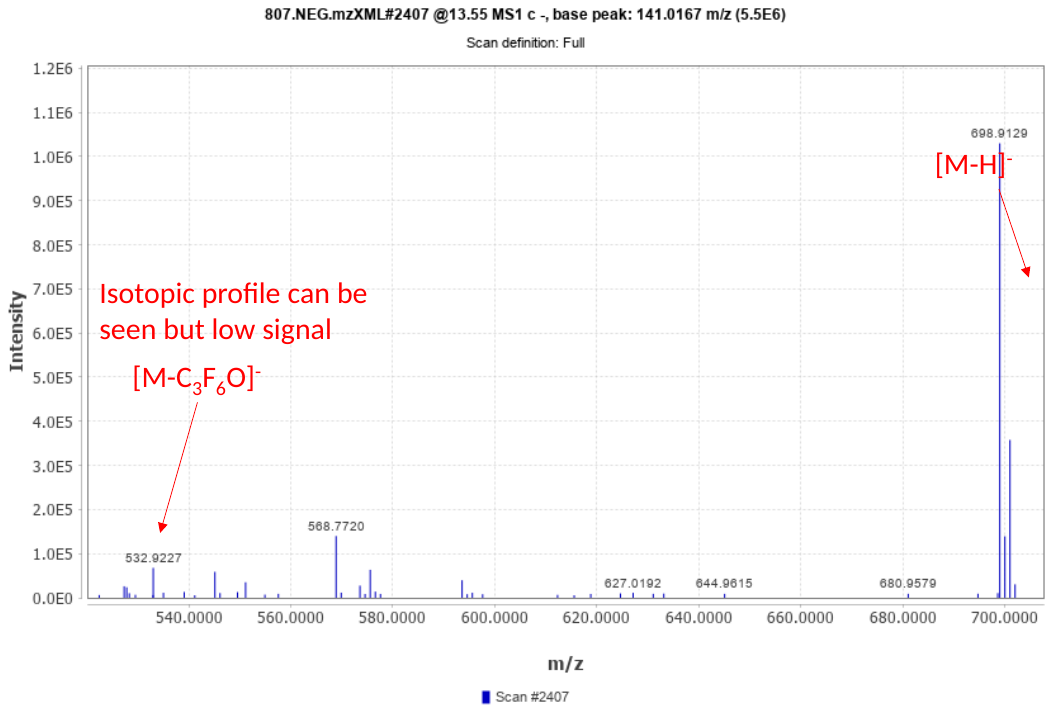

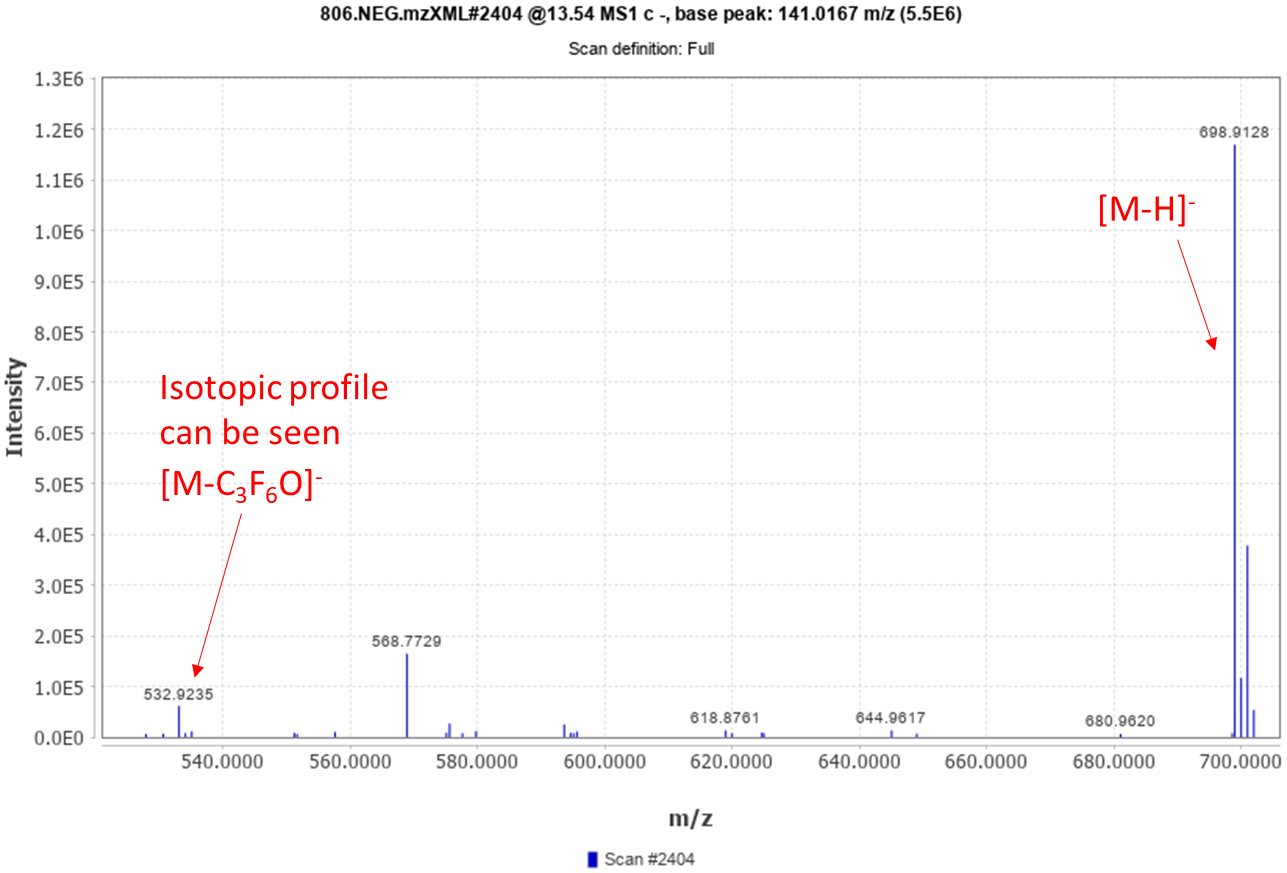

**Figure S.8.** A confirmatory fragment ([M-C_3_F_6_O]^-^) pertaining to [C_12_ClF_24_O_4_]^-^ present in MS1 from the ST001430 study and also present in the published MS/MS spectra. Mass spectra were collected from apexes of peaks using MZmine 2.0^1^.

| **Table S.8.** Comparison between calculated isotopic profile of [C_8_F_17_O_3_S]^-^ by *envi*Pat^2^ and IDSL.UFA | | | | | |
| --- | --- | --- | --- | --- | --- |
| ***envi*Pat (Centroid, ppm = 10)** | | **IDSL.UFA (peak spacing = 0.005 Da)** | | **Comparison** | |
| **Mass (Da)** | **Profile intensity (%)** | **Mass (Da)** | **Profile intensity (%)** | **Mass error (mDa)** | **Absolute intensity error (%)** |
| 498.9302176 | 100 | 498.929669 | 100 | 0.05486 | 0 |
| 499.9332550 | 9.575629 | 499.932698 | 9.491 | 0.05570 | 0.084629 |
| 500.9260979 | 4.554898 | 500.925465 | 4.602 | 0.06329 | 0.047102 |
| 500.9355769 | 0.953582 | 500.934623 | 1.016 | 0.09539 | 0.062418 |
|  |  |  |  | NEME ≤ 0.69 mDa | PCS ≥ 99.999% |

### S.2. Rdisop command details

```
if (!require("BiocManager", quietly = TRUE))
    install.packages("BiocManager")

BiocManager::install("Rdisop")
```

\## Kynurenine adduct 1 (pool_P1_A1-A12.mzML)

KynurenineAdduct1 <- matrix(c(209.09205, 86902971, 210.09537, 10.848*86902971/100), ncol = 2, byrow = TRUE)  # C10H13N2O3

Rdisop_KynurenineAdduct1_results <- decomposeIsotopes(KynurenineAdduct1[, 1], KynurenineAdduct1[, 2], ppm=10, mzabs = 0.005, elements= initializeElements(c("C", "H", "N", "O", "P", "S")), filter = NULL, z=1, maxisotopes = 2, minElements = "C2H0N0O0P0S0", maxElements = "C33H78N9O21P2S2")

\## Kynurenine adduct 2 (pool_P1_A1-A12.mzML)

KynurenineAdduct2 <- matrix(c(192.06553, 463805, 193.0689, 10.184*463805/100), ncol = 2, byrow = TRUE)  # C10H10NO3

Rdisop_KynurenineAdduct2_results <- decomposeIsotopes(KynurenineAdduct2[, 1], KynurenineAdduct2[, 2], ppm=10, mzabs = 0.005, elements= initializeElements(c("C", "H", "N", "O", "P", "S")), filter = NULL, z=1, maxisotopes = 2, minElements = "C2H0N0O0P0S0", maxElements = "C33H78N9O21P2S2")

\## PFOS (D_194NEG.mzML)

PFOS <- matrix(c(498.9312134, 162067.5625, 499.9349976, 11750.08887), ncol = 2, byrow = TRUE)  # C8F17O3S

Rdisop_PFOS_results <- decomposeIsotopes(PFOS [, 1], PFOS [, 2], ppm=10, mzabs = 0.01, elements= initializeElements(c("C", "H", "N", "F", "I","O", "P", "S")), filter = NULL, z=1, maxisotopes = 2, minElements="C4H0F0I0N0O0P0S0", maxElements="C32H77F31I4N9O10P1S1")

\## PFOS isomer 1 (1.NEG.mzXML)

PFOS_isomer1 <- matrix(c(498.92945, 100, 499.93308, 2.80, 500.92529, 0.43), ncol = 2, byrow = TRUE)  # C8F17O3S

Rdisop_PFOS1_results <- decomposeIsotopes(PFOS_isomer1 [, 1], PFOS_isomer1 [, 2], ppm=10, mzabs = 0.005, elements= initializeElements(c("C", "H", "Br", "Cl", "F", "N", "O", "P", "S")), filter = NULL, z=1, maxisotopes = 3, minElements="C2H0Br0Cl0F0N0O0P0S0",maxElements="C24H55Br6Cl8F31N3O10P1S1")

\## PFOS isomer 2 (1.NEG.mzXML)

PFOS_isomer2 <- matrix(c(498.92955, 100, 499.9329, 4.31, 500.92555, 0.86), ncol = 2, byrow = TRUE)  # C8F17O3S

Rdisop_PFOS2_results <- decomposeIsotopes(PFOS_isomer2 [, 1], PFOS_isomer2 [, 2], ppm=10, mzabs = 0.005, elements= initializeElements(c("C", "H", "Br", "Cl", "F", "N", "O", "P", "S")), filter = NULL, z=1, maxisotopes = 3, minElements="C2H0Br0Cl0F0N0O0P0S0",maxElements="C24H55Br6Cl8F31N3O10P1S1")

| **Table S.9.** Comparison of ranks of two adducts of Kynurenine (C_10_H_12_N_2_O_3_) in ”pool_P1_A1-A12.mzML” from MSV000088661 study among three different annotation methods on a simple chemical space (CHNOPS) | | | | | | | | | | | | | | |
| --- | --- | --- | --- | --- | --- | --- | --- | --- | --- | --- | --- | --- | --- | --- |
| **Rdisop** | **Peak ID** |  | **Formula Ion** | **m/z Isotopic Profile** | **m/z** | **RT * (min)** | **Peak Height** | minElements = "C2H0N0O0P0S0"  maxElements = "C33H78N9O21P2S2" | | | | | **Rank** | |
|  | 270 |  | [C_10_H_12_N_2_O_3_+H]^+^ | 209.09262 | 209.09205 | 4.167 | 1.99E+07 |  |  |  |  |  | 1 (out of 25) | |
|  | 219 |  | [C_10_H_12_N_2_O_3_+H-NH_3_]^+^ | 192.06607 | 192.06555 | 4.167 | 1.14E+07 |  |  |  |  |  | 1 (out of 17) | |
| **Enumerated chemical space (ECS)**  **(3,790,768 unique isotopic profiles)** | **Peak ID** | **ID Formula Ion** | **Formula Ion** | **m/z Isotopic Profile** | **m/z peakList** | **RT * (min)** | **Peak Height** | **NEME (mDa)** | **PCS (‰)** | **R^13^C ** peakList** | **R^13^C ** Isotopic Profile** | **NDCS** | **RCS (%)** | **Rank** |
|  | 270 | 692589 | [C_10_H_12_N_2_O_3_+H]^+^ | 209.09262 | 209.09205 | 4.167 | 1.99E+07 | 0.58 | 1000 | 10.85 | 10.71 | 5 | 100 | 1 |
|  | 219 | 689281 | [C_10_H_12_N_2_O_3_+H-NH_3_]^+^ | 192.06607 | 192.06555 | 4.167 | 1.14E+07 | 0.54 | 1000 | 10.89 | 10.71 | 5 | 100 | 1 |
| **IDSL.ExposomeDB**  **(65,133 unique isotopic profiles)** | **Peak ID** | **ID Formula Ion** | **Formula Ion** | **m/z Isotopic Profile** | **m/z peakList** | **RT * (min)** | **Peak Height** | **NEME (mDa)** | **PCS (‰)** | **R^13^C ** peakList** | **R^13^C ** Isotopic Profile** | **NDCS** | **RCS (%)** | **Rank** |
|  | 270 | 445 | [C_10_H_12_N_2_O_3_+H]^+^ | 209.09262 | 209.09205 | 4.167 | 1.99E+07 | 0.61 | 1000 | 10.85 | 11.02 | 5 | 100 | 1 |
|  | 219 | 340 | [C_10_H_12_N_2_O_3_+H-NH_3_]^+^ | 192.06607 | 192.06555 | 4.167 | 1.14E+07 | 0.57 | 1000 | 10.89 | 10.97 | 5 | 100 | 1 |
| * **RT** (retention time)  ** **R^13^C** = ratio of intensity of ^13^C isotopologue to ^12^C isotopologue (%) | | | | | | | | | | | | | | |

| **Table S.10.** Comparison of ranks of two isomers of PFOS ([C_8_F_17_O_3_S]^-^) in”1.NEG.mzXML” from ST001430 study between annotation methods on a complex chemical space ( CHBrClFNOPS) | | | | | | | | | | | | | | | | | | | | | | |
| --- | --- | --- | --- | --- | --- | --- | --- | --- | --- | --- | --- | --- | --- | --- | --- | --- | --- | --- | --- | --- | --- | --- |
| **Rdisop** | **Peak ID** |  | **Formula Ion** | | **m/z Isotopic Profile** | | | **m/z** | | **RT * (min)** | | **Peak Height** | | minElements="C2H0Br0Cl0F0N0O0P0S0"  maxElements="C24H55Br6Cl8F31N3O10P1S1" | | | | | | | | **Rank** |
|  | 3692 |  | [C_8_F_17_O_3_S]^-^ | | 498.92967 | | | 498.92962 | | 10.292 | | 3.35E+05 | |  |  |  |  |  |  |  |  | 31 ^†^ |
|  | 3693 |  | [C_8_F_17_O_3_S]^-^ | | 498.92967 | | | 498.92953 | | 10.537 | | 1.07E+06 | |  |  |  |  |  |  |  |  | 22 ^††^ |
| ^†^ 31 out of 3461 candidate molecular formulas  ^††^ 22 out of 3459 candidate molecular formulas | | | | | | | | | | | | | | | | | | | | | | |
| **Enumerated chemical space (ECS)**  **(7,135,620 unique isotopic profiles)** | **Peak ID** | **ID Formula Ion** | | **Formula Ion** | | **m/z Isotopic Profile** | | | **m/z peakList** | | **RT * (min)** | | **Peak Height** | | **NEME (mDa)** | | **PCS (‰)** | **R^13^C ** peakList** | **R^13^C ** Isotopic Profile** | **NDCS** | **RCS (%)** | **Rank** |
|  | 3692 | 428596 | | [C_8_F_17_O_3_S]^-^ | | 498.92967 | | | 498.92945 | | 10.292 | | 3.35E+05 | | 0.16 | | 998 | 2.86 | 8.57 | 13 | 36.11 | 1 |
|  | 3693 | 428596 | | [C_8_F_17_O_3_S]^-^ | | 498.92967 | | | 498.92955 | | 10.537 | | 1.07E+06 | | 0.12 | | 999 | 4.41 | 8.57 | 14 | 58.33 | 1 |
| **IDSL.ExposomeDB**  **(129,122 unique isotopic profiles)** | **Peak ID** | **ID Formula Ion** | | **Formula Ion** | | **m/z Isotopic Profile** | | | **m/z peakList** | | **RT * (min)** | | **Peak Height** | | **NEME (mDa)** | | **PCS (‰)** | **R^13^C ** peakList** | **R^13^C ** Isotopic Profile** | **NDCS** | **RCS (%)** | **Rank** |
|  | 3692 | 39056 | | [C_8_F_17_O_3_S]^-^ | | 498.92967 | | | 498.92945 | | 10.292 | | 3.35E+05 | | 0.16 | | 998 | 2.86 | 8.69 | 13 | 36.11 | 1 |
|  | 3693 | 39056 | | [C_8_F_17_O_3_S]^-^ | | 498.92967 | | | 498.92955 | | 10.537 | | 1.07E+06 | | 0.13 | | 999 | 4.41 | 8.69 | 14 | 58.33 | 1 |
| * **RT** (retention time)  ** **R^13^C** = ratio of intensity of ^13^C isotopologue to ^12^C isotopologue (%) | | | | | | | | | | | | | | | | | | | | | | |
| 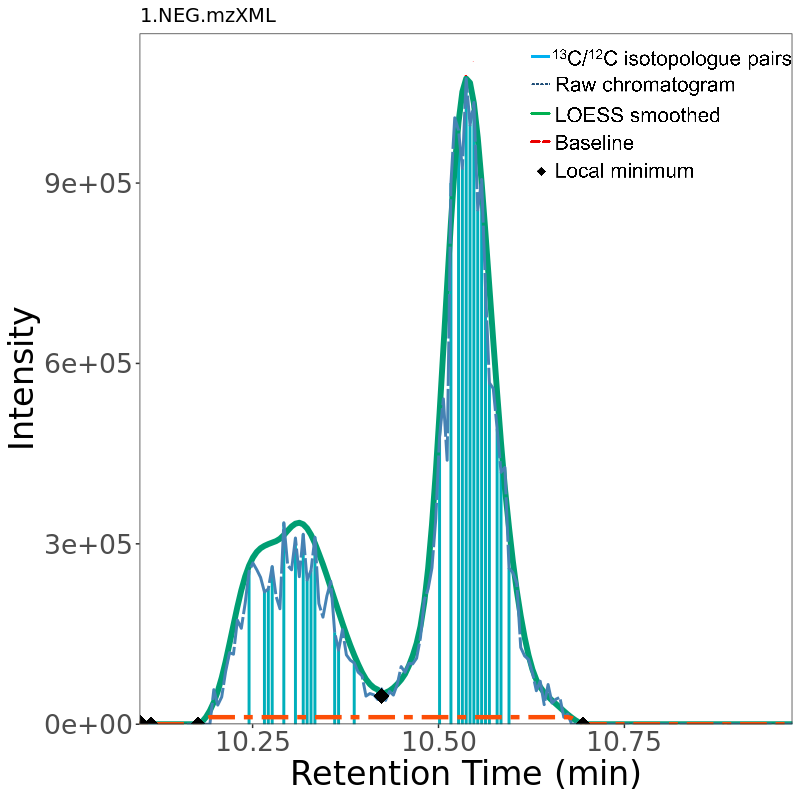 | | | | | | | 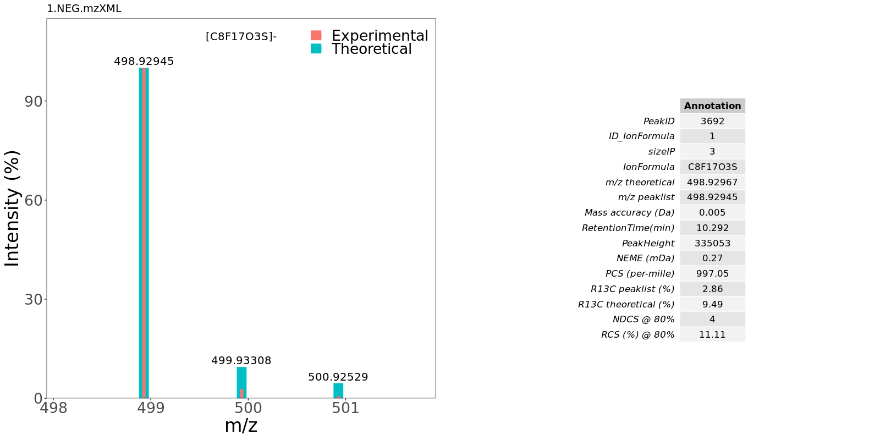RT=10.292 | | | | | | | | | 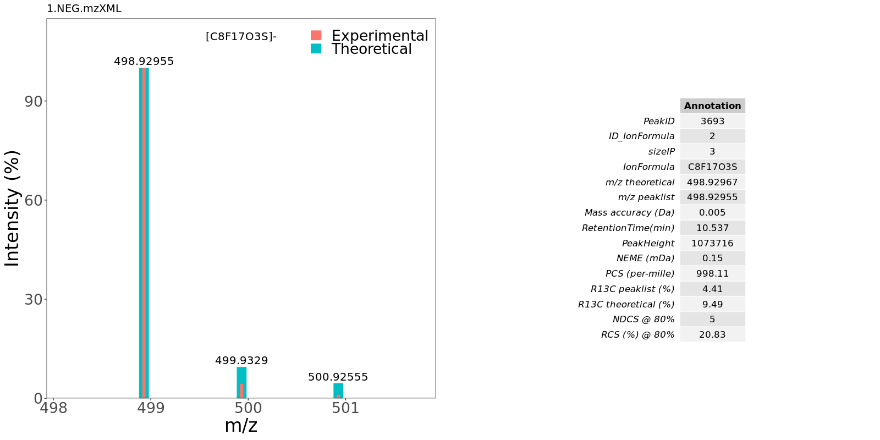RT=10.537 | | | | | | |
| **Figure S.9.** EIC and integrated spectra of the PFOS peaks in 1.NEG.mzXML” from ST001430 | | | | | | | | | | | | | | | | | | | | | | |

| **Table S.11.** Comparison of ranks of PFOS (C_8_HF_17_O_3_S) in ”D_194NEG.mzML” from ST002044 study between annotation methods on a complex chemical space ( CHBrClFNOPS) | | | | | | | | | | | | | | | |
| --- | --- | --- | --- | --- | --- | --- | --- | --- | --- | --- | --- | --- | --- | --- | --- |
| **Rdisop** | **Peak ID** |  | **Formula Ion** | **m/z Isotopic Profile** | **m/z** | **RT * (min)** | | **Peak Height** | minElements="C4H0F0I0N0O0P0S0"  maxElements="C32H77F31I4N9O10P1S1" | | | | | **Rank** | |
|  | 6675 |  | [C_8_F_17_O_3_S]^-^ | 498.92967 | 498.9312134 | 9.982 | | 1.78E+05 |  |  |  |  |  | 36 (out of 1095) | |
| **IDSL.ExposomeDB**  **(129,122 unique isotopic profiles)** | **Peak ID** | **ID Formula Ion** | **Formula Ion** | **m/z Isotopic Profile** | **m/z peakList** | **RT * (min)** | | **Peak Height** | **NEME (mDa)** | **PCS (‰)** | **R^13^C ** peakList** | **R^13^C ** Isotopic Profile** | **NDCS** | **RCS (%)** | **Rank** |
|  | 6675 | 39056 | [C_8_F_17_O_3_S]^-^ | 498.92967 | 498.93084 | 9.982 | | 1.78E+05 | 1.95 | 998 | 2.79 | 9.49 | 3 | 37.5 | 1 |
| * **RT** (retention time)  ** **R^13^C** = ratio of intensity of ^13^C isotopologue to ^12^C isotopologue (%) | | | | | | | | | | | | | | | |
| 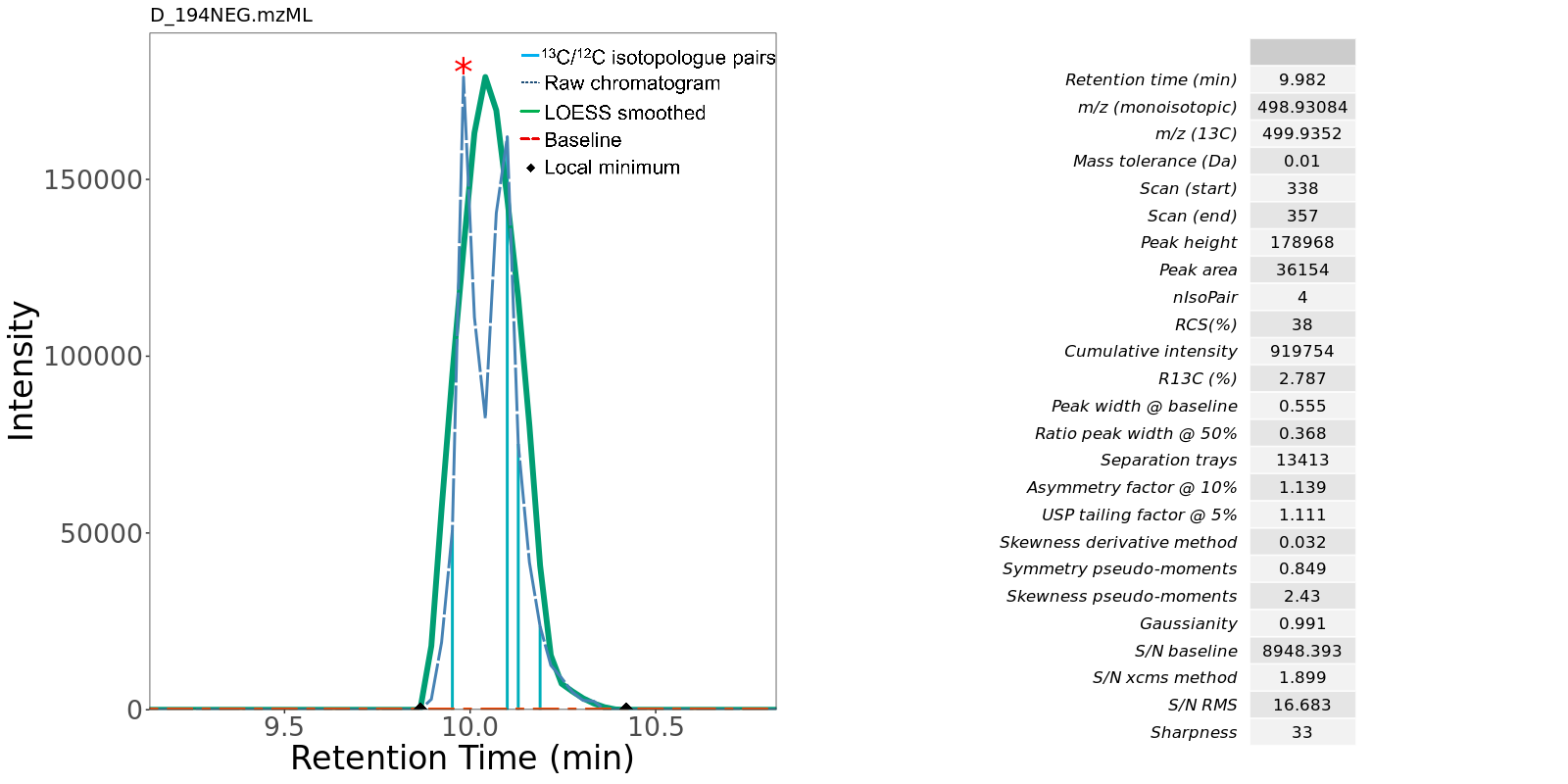 | | | | | | | 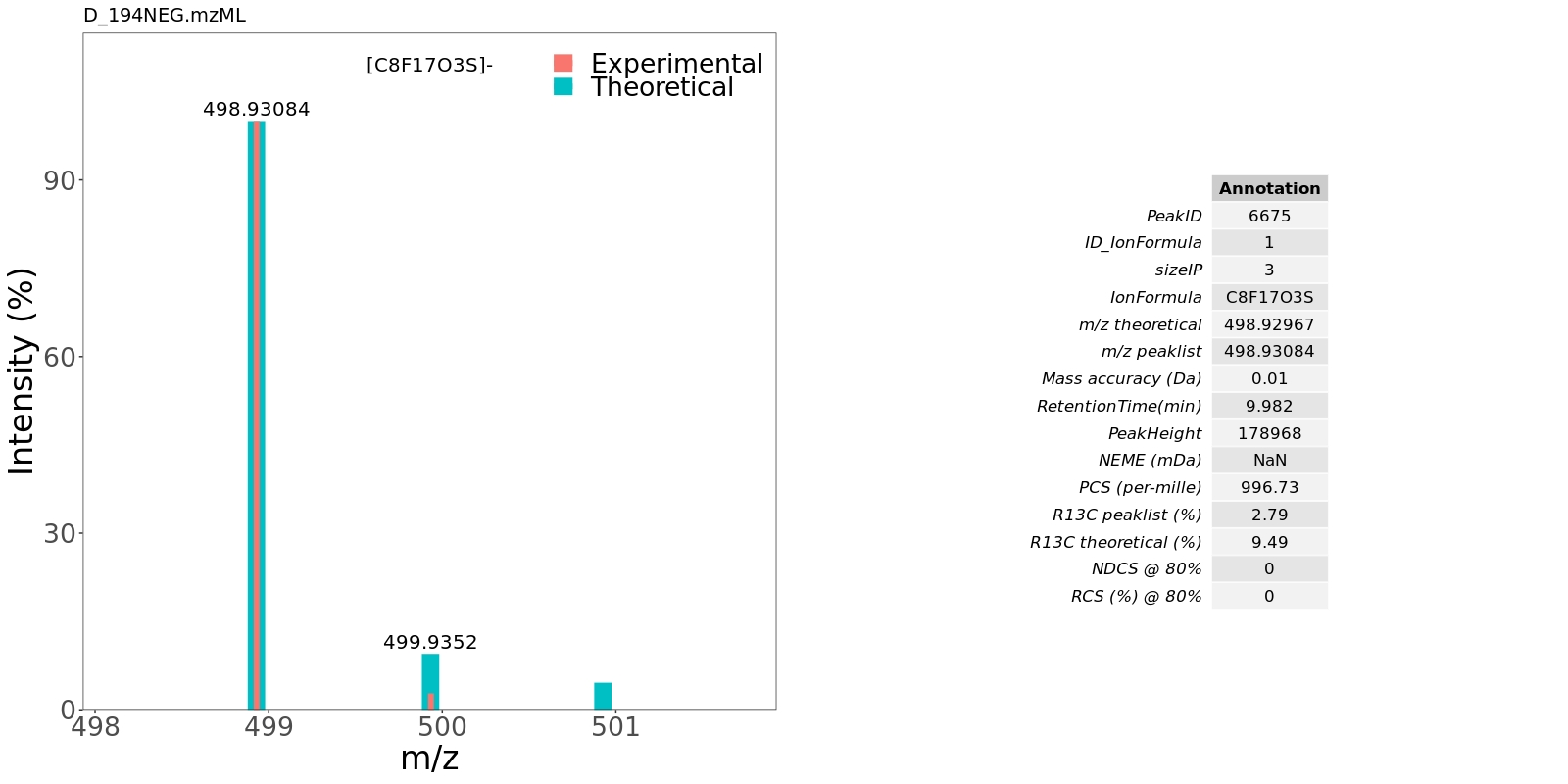 | | | | | | | | |
| **Figure S.10.** EIC and integrated spectra of the PFOS peak in ”D_194NEG.mzML” from ST002044 study | | | | | | | | | | | | | | | |

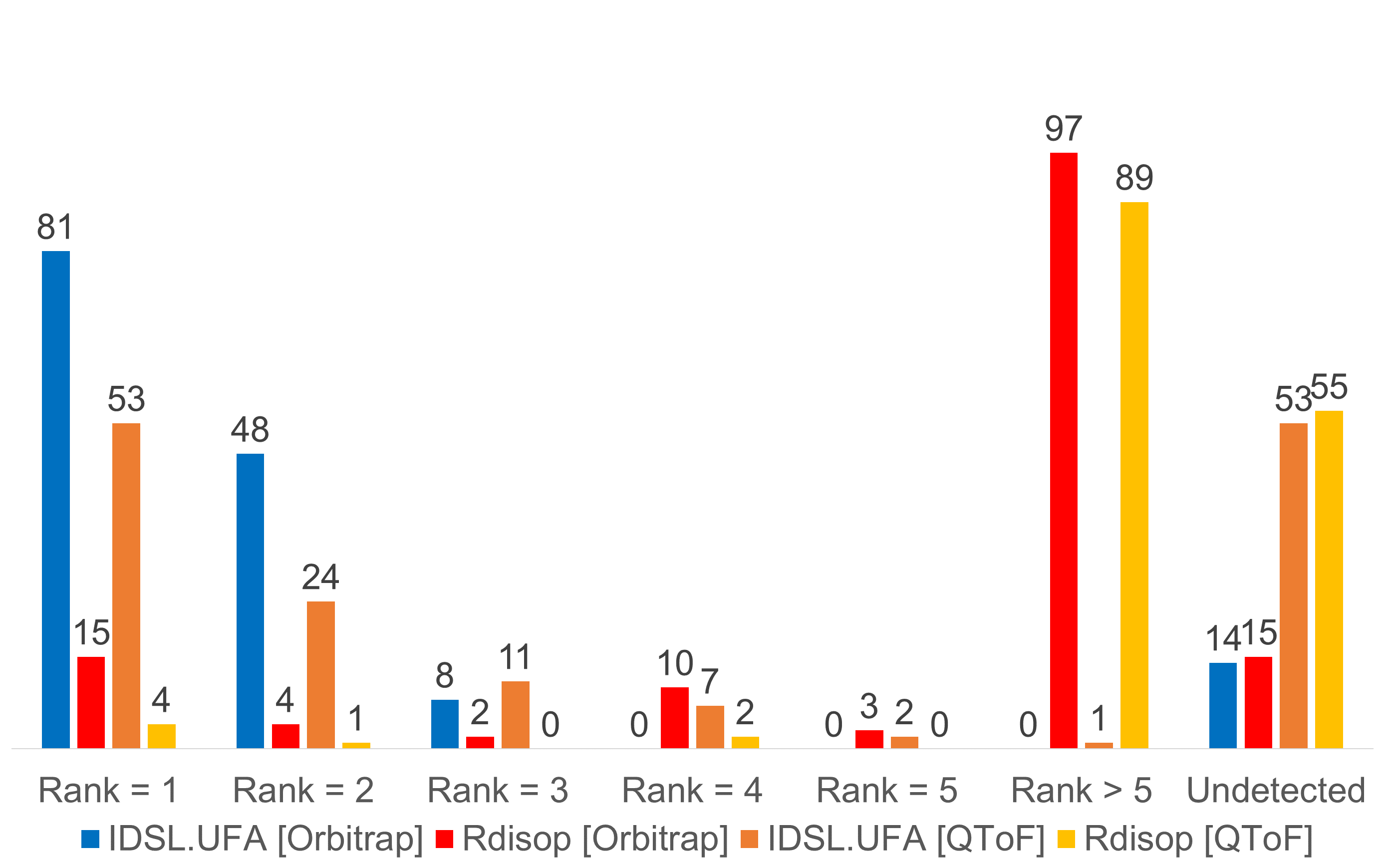

**Figure S.11.** Comparison of ranks of 151 true positive hits between IDSL.UFA and Rdisop. Orbitrap Data were from file “021518_387057_CSHn_BioRec2.mzML” from ST001264 and QTOF data were from “Biorec005_negCSH_postKieffer040.mzML” from ST001843 study. True positive annotations were from study ST001154. All three datasets were generated by same chromatography method as reported in <https://www.metabolomicsworkbench.org/data/DRCCMetadata.php?Mode=Study&StudyID=ST001154>

| **Table S.12.** Summarized operational comparison between IDSL.UFA and Rdisop^3^ | | |
| --- | --- | --- |
| **Criteria** | **IDSL.UFA** | **Rdisop^3^** |
| Can annotate a LC/HRMS data file with formulas? | Yes | Yes, but need additional scripting |
| Query the raw data for a theoretical isotope profile? | Yes | Yes, but need additional scripting |
| Can handle more than 6 elements for predicting formula for a single m/z? | Yes, up to 16 | Yes |
| Can batch process an entire study? | Yes | No |
| Generate ranking using the aligned peak table? | Yes | Yes, but need additional scripting |
| Provide scores to rank formula hits for an experimental isotope profile? | Yes | Yes |
| Can compute isotope profile from a single formula? | Yes | Yes |
| Can compute isotope profiles for a list of formula from a database? | Yes | Yes, but need additional scripting |
| Can compute formula for common ESI adducts? | Yes | Yes |
| Can provide formulas for a single m/z value? | Yes | Yes |
| Can predict elemental composition from the experimental isotope profile? | Yes | Yes |
